# Signal Associa: hot tumors elicit field-spanning NK cell coordination linked to their recruitment

**DOI:** 10.64898/2026.10.21.760734

**Authors:** Majid Rostami, Hyeri Choi, Yeonwoo Rho, Noo Li Jeon, Sangyoon J. Han

## Abstract

Cells coordinate through coupled dynamics, yet whether observed correlations reflect genuine coupling is rarely tested. Signal Associa is an open-source framework uniting six coupling methods with automated stationarization and surrogate testing. Benchmarking against six ground truth systems reveals a trichotomy of coupling regimes, each with a different best method: correlation for coupling carried by shared trends and cycles, Granger causality for linear stochastic coupling, and convergent cross-mapping for nonlinear coupling; agreement across methods identifies the regime from data. In 13 movies of natural killer (NK) cells across three colorectal tumor microenvironments, NK cells formed field-spanning coupling networks at a conserved rate; at matched exposure to tumor signal, NK cells around hot tumors were more globally integrated, and hot tumors with a more integrated NK network tended to attract more NK cells. In focal adhesions, stiffness strengthened vinculin–FAK coupling, most at protruding edges. Surrogates cut lagged-correlation false positives from 23–92% to about 5%.

## Introduction

Quantifying coupling between simultaneously recorded time series is a foundational problem across the biological sciences. In neuroscience, coupling analysis between neural spike trains and calcium transients reveals functional connectivity and information flow in circuits [1, 2]. In ecology, coupling between species abundance time series identifies predator–prey interactions and shared environmental drivers [3]. In cell biology, coupling between fluorescence intensity traces at subcellular structures exposes signaling hierarchies and feedback regulation [4]. In gene regulatory network inference, coupling measures applied to expression time series reconstruct causal regulatory relationships [5]. Across all these domains, the central question is the same: given two signals that co-vary in time, are they truly coupled, and how can we validate the answer?

The challenge is that biological coupling does not occupy a single regime. In the simplest case, coupling propagates through stochastic fluctuations, where the past of one signal linearly predicts the future of another; this is the regime standard methods implicitly assume. But many biological systems depart from it in ways that invalidate those methods. Nonlinear dynamical coupling, as in interacting biochemical oscillators [6], produces time series whose long-term correlation averages to zero despite genuine causal interaction, because the relationship is state-dependent and alternates in sign [3]. In molecular-clutch mechanotransduction, where the traction response is biphasic in substrate stiffness [7] and therefore is set by magnitude instead of sign, non-monotonic coupling forms, which violates the linear correlation assumption. Shared external inputs, such as a common upstream regulator driving two downstream targets, create apparent coupling where none exists [8]. Both deterministic trends and cycles, as in focal-adhesion assembly and disassembly [9], and imaging artifacts such as photobleaching [10] or instrument drift, introduce non-stationarity that violates the stationarity assumption underlying classical statistical inference. Given that any of these coupling mechanisms may be present, the central question is which method is most pertinent to a given inference problem, so as to avoid both missing true coupling and reporting false positives.

In practice, most biological coupling studies rely on Pearson or Spearman correlation. Correlation is instantaneous and symmetric: it compares *X*(*t*) to *Y* (*t*) as isolated pairs of values, with no memory of past dynamics and no ability to distinguish cause from effect. A system where *Y* (*t*) = *f* (*X*(*t*−5)) with a five-step delay yields *r* ≈ 0. Lag-scanning [11], introduced for Rho GTPase coupling analysis, addresses this issue and is widely adopted since [12, 13], remaining the standard in cell biology. Despite its strength in capturing co-variation across different types of signals, it faces two fundamental barriers: it cannot detect the direction of influence, whether unilateral or bilateral; and it is unable to capture nonlinear, non-monotonic, or state-dependent coupling, where the relationship is sign-symmetric or alternates in sign: correlation remains near zero regardless of lag.

For studies requiring directionality, Granger causality has emerged as the leading alternative [14, 15]. But applying GC carries a fundamentally flawed assumption: it requires stationarity [16, 17] and assumes that biologically relevant coupling resides exclusively in the stochastic component of the raw signal. In practice, trends, oscillations, and other slow deterministic components must be removed before GC can be applied. Lai et al. [14] manually subtracted low-frequency components from actin regulator signals before GC, discarding them without testing whether they contained biologically relevant coupling. Even pipelines that advance beyond classical GC inherit this assumption: CausalXtract [15], a causal-discovery framework for live-cell imaging, assumes the dynamics are stationary and offers no way to detect or handle non-stationary components. Ravishankar et al. [13] explicitly rejected GC and transfer entropy for Rho GTPase signaling analysis, citing stationarity sensitivity, and retreated to Pearson. Yet in systems where the coupling is mediated by an oscillating value or through feedback loops between two nonlinear signals, the coupling resides precisely in the deterministic components that stationarization destroys.

Once a method detects coupling between two signals, a second problem concerns how its validity can be assessed against chance. Commonly, parametric tests are used, which assume independent samples; autocorrelation between biological signals violates this assumption [18] and, as we show in the Results, can induce spurious coupling detection and thereby high false-positive rates. Alternatively, surrogate-based testing avoids this by generating randomized signals that preserve each signal’s spectral and distributional properties while destroying cross-signal coupling [19]. The initially detected coupling must survive this structured null to be labeled genuine.

Here we present Signal Associa, an open-source framework for regime-aware biological coupling analysis. The core insight is that no single method is valid across all coupling regimes: correlation (Pearson and Spearman) captures linear co-variation but is blind to directionality and nonlinearity; Granger causality provides directionality but requires stationarity and destroys deterministic coupling; and convergent cross-mapping detects nonlinear, state-dependent, and deterministic coupling but is more conservative than Granger causality in purely linear regimes. Rather than choosing one method, Signal Associa applies all three classes in parallel, together with complementary measures (mutual information and transfer entropy), with automated stationarization, surrogate-based significance testing, and GPU-accelerated computation, letting the data reveal which regime each signal pair occupies. We validate this approach against six ground-truth systems spanning linear, nonlinear, non-monotonic, deterministic, and time-varying coupling, together with five sensitivity experiments that probe robustness to surrogate choice, signal length, observation noise, lag scanning, and shared drivers.

We apply Signal Associa to two biological systems in which the coupling between simultaneously imaged signals may be missed by linear methods. In the first application, we ask whether natural killer cells coordinate their migration during tumor immunosurveillance, and whether the tumor microenvironment modulates this coordination, by analyzing trajectory coupling across three molecularly distinct tumor types. In the second, we ask how substrate stiffness reshapes the coupling between vinculin recruitment and FAK phosphorylation at individual focal adhesions.

## Results

### The Signal Associa framework

Signal Associa is a modular Python framework that unifies stationarization, surrogate generation, and six association methods into a single pipeline for analyzing coupling between pairs of biological time series (Fig 1a). The pipeline runs in five stages: (i) input, taking the two raw signals to be tested (Fig 1b); (ii) stationarization, which removes trends and cycles that can produce spurious correlations by decomposing each signal as *X*(*t*) = *T* (*t*) + *C*(*t*) + *S*(*t*) + *ε*(*t*) and stripping the deterministic components, with success judged by a battery of stationarity diagnostics—the augmented Dickey–Fuller (ADF) and KPSS tests, the Phillips–Perron test, a rolling-variance ratio, and spectral concentration (Fig 1c; Fig S6); (iii) method routing, which sends the stationarized signal to Granger causality and the raw signals to the other five methods, because only Granger causality requires stationarity; (iv) surrogate significance testing, which builds a null by generating *N* synthetic signals that keep each signal’s statistical properties but break any coupling (IAAFT, shuffle, circular shift, or block bootstrap) and reports an empirical p-value from where the observed statistic falls in that null (Fig 1d–e); and (v) a multi-method summary showing which methods agree and disagree on coupling (Fig 1e). Full algorithmic details for each stage are provided in Materials and methods; mathematical derivations and implementation notes are in S1–S7.

**Fig 1.**
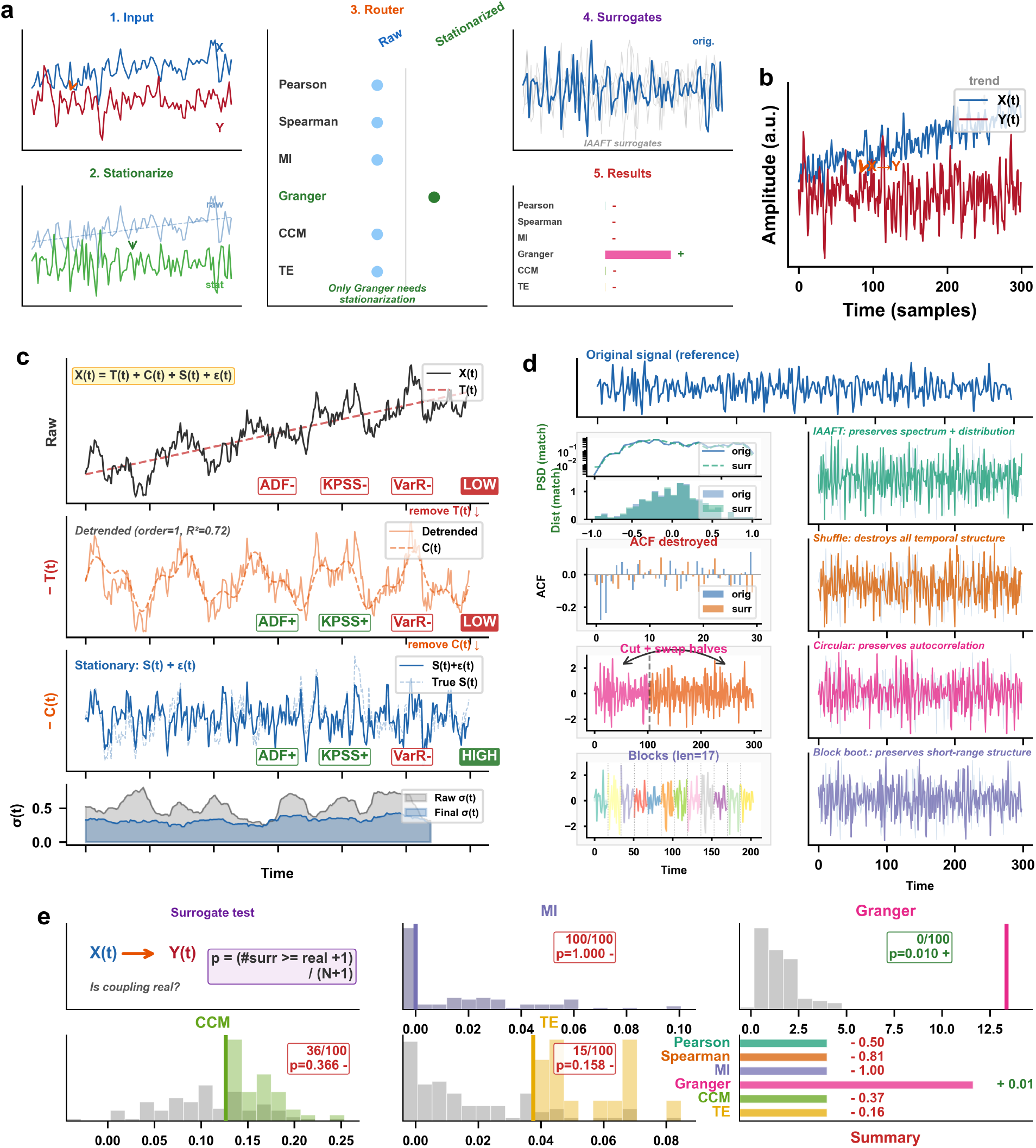
The Signal Associa pipeline for unified coupling analysis. (**a**) Five-stage pipeline overview: input signals, stationarization, method routing (raw vs stationary), surrogate generation, and results; panels (**b–e**) expand these stages in turn on a worked example. (**b**) Example coupled autoregressive AR(2) input with unidirectional *X* → *Y* coupling and linear trend on *X*. (**c**) Stationarization decomposing *X*(*t*) = *T* (*t*)+*C*(*t*)+*S*(*t*)+*ε*(*t*): trend removal, cycle removal, and stationarity validation via ADF/KPSS/variance ratio diagnostics at each step. (**d**) Four surrogate methods with mechanism insets: IAAFT (preserves spectrum and distribution), shuffle (destroys temporal structure), circular shift (preserves autocorrelation), and block bootstrap (preserves short-range structure). (**e**) Surrogate significance testing: null distributions from 100 IAAFT surrogates (gray) with real statistics (colored lines) for MI, Granger, CCM, and TE. Bottom-right: summary of all six methods with p-values and significance decisions.

This multi-method view is the core design principle: rather than relying on a single coupling measure, Signal Associa presents all six methods simultaneously, enabling researchers to distinguish linear from nonlinear coupling, assess directionality, and identify potential confounders.

### Ground truth benchmarking

To evaluate the sensitivity and specificity of each association method, and to characterize how surrogation and stationarization affect detection performance, we tested Signal Associa against six ground truth systems with known coupling structures, complemented by five sensitivity experiments (surrogate choice, signal length, observation noise, lag scanning, and shared drivers). All significance testing used IAAFT surrogates (*N* = 1,000) unless otherwise stated.

#### Detection across coupling regimes

We tested all six methods on six ground truth systems spanning the coupling regimes defined in Materials and methods (100 independent pairs per coupling strength, *α* from 0 to 0.8, or up to 8.0 for the clutch system; 1,000 IAAFT surrogates; Fig 2).

**Fig 2.**
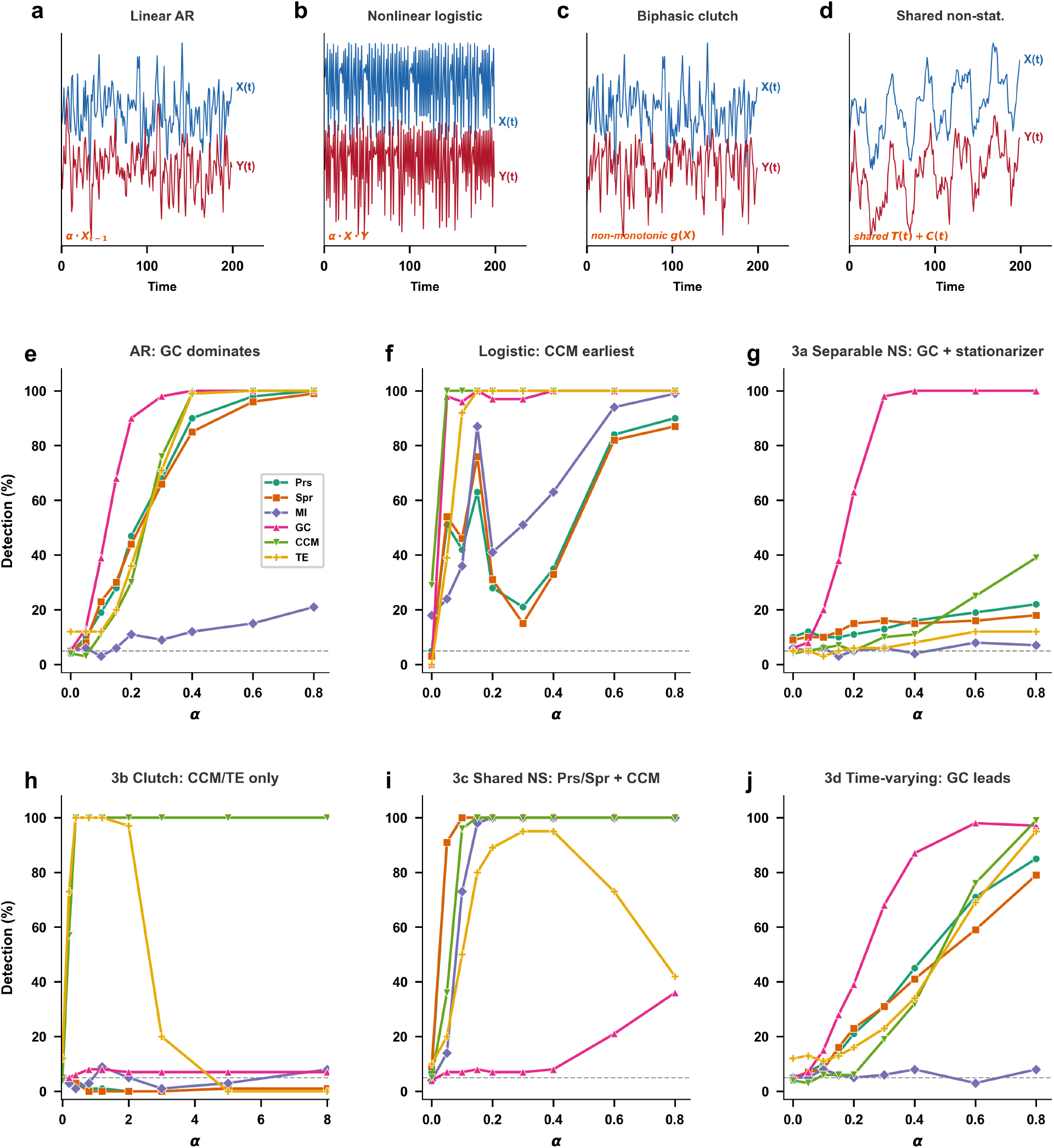
Detection across coupling regimes. (**a–d**) Representative signal pairs (*X* upper, *Y* lower): linear AR (a), nonlinear logistic (b), biphasic clutch (c; non-monotonic Bell-model coupling), and shared non-stationarity (d; coupling carried by shared trends/cycles). (**e–j**) Detection rate versus coupling strength *α* for each ground-truth system (100 pairs, 1000 IAAFT surrogates, *n* = 300 except the logistic system at *n* = 100; dashed line marks the 5% level). (**e**) AR: GC dominates, with CCM/TE following and Pearson/Spearman slower. (**f** ) Logistic: CCM and GC reach 100% earliest; Pearson/Spearman weaker (mirage correlations). (**g**) 3a Separable non-stationarity: GC recovers 100% after stationarization; all raw-signal methods fail. (**h**) 3b Biphasic clutch: CCM detects across the full range while TE detects only transiently before collapsing; Pearson/Spearman/GC stay at chance. (**i**) 3c Shared non-stationarity: Pearson/Spearman and CCM reach 100% on raw signals; GC blind, as stationarization removes the coupling. (**j**) 3d Time-varying coupling: GC leads; CCM/TE weaker as coupling is intermittent.

##### (1) Linear coupling (AR)

On coupled AR(2) processes (*n* = 300; Fig 2e), Granger causality detected coupling earliest (100% by *α* = 0.4; 98% at *α* = 0.3), consistent with its design for linear autoregressive systems. CCM and TE were more conservative, rising later than Granger causality but still reaching 99% by *α* = 0.4. Pearson and Spearman were slower to detect coupling, at 90% and 85% at *α* = 0.4 before converging to 100% and 99% by *α* = 0.8. MI showed weak sensitivity throughout (≤21%), reflecting kNN estimation limits at *n* = 300. All methods maintained false-positive rates ≤5% except TE (12%; Fig 2e).

##### (2) Nonlinear coupling (bidirectional chaotic logistic maps)

On bidirectional coupled logistic maps (Sugihara et al. [3] Eq. 1; *r*_*X*_ = 3.8, *r*_*Y*_ = 3.5, *β*_*XY*_ = 0.02 fixed, *n* = 100; Fig 2f), CCM reached 100% surrogate-based detection from *β*_*Y X*_ = 0.05, the earliest of all methods, consistent with its design for state-space reconstruction of nonlinear attractors. GC and TE followed closely, both saturating by *β*_*Y X*_ = 0.15. Pearson and Spearman showed a distinctive non-monotonic pattern: surrogate detection dropped at intermediate coupling strengths before recovering at high *β*_*Y X*_, reflecting the “mirage correlations” phenomenon [3] in which alternating windows of positive and negative correlation cancel the global statistic even when dynamical coupling is strong. MI lagged other methods throughout and showed a similar intermediate dip

##### (3a) Separable non-coupled non-stationarity

Adding deterministic trends and multi-frequency cycles to the observed driver signal (while the coupling itself remained in the underlying AR process) destroyed detection for all raw-signal methods (Pearson 16%, Spearman 15%, CCM 11%, TE 8% at *α* = 0.4; Fig 2g), as the dominant trend and cycle variance masked the coupling signal. MI remained at chance throughout. These methods thus produce false negatives, missing the autoregressive coupling beneath the non-stationary envelope. Only GC, operating on stationarized signals, recovered 100% detection from *α* = 0.3, identifying it as the most reliable method when the coupling resides in the stationary component and the non-stationarity is additive and non-coupled

##### (3b) Biphasic clutch coupling (non-monotonic)

Under non-monotonic bell-shaped coupling (Eq. (8); *v*_0_ = 0.07, *n* = 300; Fig 2h), CCM was the only method to reliably detect coupling across the full range, reaching 100% from *α* = 0.4 onward. Pearson, Spearman, MI, and GC all remained at or near chance (≤8%), as the even symmetry of the clutch function (*g*(*x*) = *g*(−*x*)) destroys any monotonic or linear relationship. TE detected coupling transiently (100% for 0.4 ≤ *α* ≤ 2.0) but collapsed to 0% at *α* ≥ 5.0, where the biphasic function saturates and the conditional entropy signal vanishes. CCM’s attractor-based reconstruction is immune to this saturation effect, demonstrating that only state-space reconstruction recovers non-monotonic coupling

##### (3c) Shared non-stationarity

When coupling resided entirely in shared deterministic trends and cycles, with independent stochastic AR dynamics, the roles reversed (Fig 2i): Pearson and Spearman reached 100% detection from *α* = 0.10, and CCM from *α* = 0.15, all operating on raw signals. GC remained at chance (≤8% up to *α* = 0.4), because stationarization removed the very components carrying the coupling. TE degraded at high coupling (95% at *α* = 0.4 but 42% at *α* = 0.8), suggesting that strong deterministic components overwhelm conditional entropy estimation. This result demonstrates that stationarization, while essential for GC, discards deterministic coupling structure that may be the biologically relevant signal.

##### (3d) Time-varying coupling strength

When coupling strength oscillated sinusoidally (*α*(*t*) = *α*_0_(1 + sin(2*πt/P* ))*/*2, *P* = 100, *n* = 300; Fig 2j), GC was the strongest detector, reaching 87% surrogate-based detection at *α*_0_ = 0.40 and 98% at *α*_0_ = 0.60. Although the coupling varies over time, the relationship remains linear and both signals are individually stationary, so the VAR model captures the average linear dependence; unlike the shared-non-stationarity case, stationarization here removes nothing that carries the coupling. Pearson and Spearman tracked each other closely but lagged GC throughout (45% and 41% at *α*_0_ = 0.40; 85% and 79% at *α*_0_ = 0.80), as the global correlation averages over coupled and uncoupled windows, diluting the effective signal. CCM lagged further, reaching 76% at *α*_0_ = 0.60 and 99% only at *α*_0_ = 0.80: the uncoupled half-periods contribute manifold regions where *Y* does not encode *X*, diluting the cross-map correlation. TE followed CCM closely. MI remained at chance throughout (≤8%)

#### Robustness analysis

##### (1) Surrogate testing versus parametric significance

A critical methodological finding emerged from applying lag-scanned Pearson correlation [11] with parametric significance across the six ground-truth systems at *α* = 0 (no coupling). Following the original method, significance was assessed at the 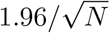 confidence level for Pearson’s coefficient, which assumes *N* independent observations, a condition violated by autocorrelation in biological time series [18]. Under this criterion, false-positive rates spanned 23–92% across the six systems, highest in the most strongly autocorrelated systems (Table 1). IAAFT surrogate testing, by contrast, controlled the false-positive rate at or below 5% in all systems, with the parametric inflation scaling directly with the autocorrelation the 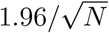 rule ignores: strongly autocorrelated systems (AR, clutch, time-varying) reached 92%, while the near-white logistic map showed the least (23%). These results identify surrogate testing as essential and IAAFT as a reliable null model, motivating Signal Associa’s design decision to report surrogate-based *p*-values as the primary significance measure for all methods

##### (2) Lag scanning does not rescue correlation for nonlinear coupling

Given the failure of classic zero-lag Pearson for non-monotonic and nonlinear signals, the lag-scanned Pearson correlation [11], the standard practice in the field, might seem able to recover missed couplings. We therefore applied the full lag-scanned procedure (peak |*ρ*| over lags [−15, +15], IAAFT significance) to every GT system at moderate coupling (*α* = 0.4) and compared it against classic lag-0 Pearson and Spearman under the same surrogate test (Table 2). Lag scanning markedly improved AR and especially the logistic case but still failed on the biphasic clutch (4%, identical to lag-0), where the even, non-monotonic coupling function *g*(*x*) = *g*(−*x*) produces near-zero linear correlation at every lag. It recovered separable non-stationarity and time-varying coupling only partially (32% and 67%). Together, lag scanning extends correlation’s reach in time but remains non-functional when the mean is not a pertinent representative of the signal, as in the biphasic case: the mean-centered products 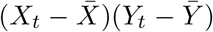 that define *ρ* (Eq. 21) cancel across positive and negative deviations, leaving *ρ* ≈ 0 at every lag.

**Table 1.** Lag-scanned Pearson false positive rates at *α* = 0 (no coupling). = 1.96 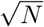 confidence level [11]; IAAFT = surrogate-based *p*-value. 1000 surrogates, 100 pairs per system.

| GT System | Parametric FPR | IAAFT FPR |
| --- | --- | --- |
| (1) AR | 92% | 3% |
| (2) Logistic | 23% | 5% |
| (3a) Separable NS | 72% | 3% |
| (3b) Clutch | 92% | 3% |
| (3c) Shared NS | 72% | 3% |
| (3d) Time-varying | 92% | 3% |

**Table 2.** **Detection at** *α* = 0.4 **(moderate coupling): lag-scanned versus classic lag-0 Pearson/Spearman**, all with IAAFT surrogate significance. 1000 surrogates, 100 pairs per system.

| GT System | Lag-scanned Prs | Classic Prs | Classic Spr |
| --- | --- | --- | --- |
| (1) AR | 100% | 90% | 85% |
| (2) Logistic | 100% | 35% | 33% |
| (3a) Separable NS | 32% | 16% | 15% |
| (3b) Clutch | 4% | 4% | 3% |
| (3c) Shared NS | 100% | 100% | 100% |
| (3d) Time-varying | 67% | 45% | 41% |

##### (3) Different types of surrogation and randomization

We compared IAAFT, shuffle, block bootstrap, and circular surrogates across three GT systems (linear AR, *α* = 0.30; biphasic clutch, *α* = 1.20; and shared non-stationarity, *α* = 0.10) at both null (*α* = 0.0) and coupled strengths (Fig 3g,h). GC was surrogate-insensitive across every condition (3–5% at null, unchanged detection when coupled). CCM was the most surrogate-dependent: block bootstrap collapsed CCM detection from 76% to 1% on AR and from 96% to 8% on shared non-stationarity, because preserving local temporal blocks retains the manifold structure that cross-mapping exploits, destroying the contrast against the surrogate null. Shuffle inflated false positives on the shared non-stationarity null (Pearson/Spearman 7–9% → 18–19%, CCM 5% → 14%) by breaking the autocorrelation that IAAFT preserves. IAAFT and circular surrogates produced comparable results throughout, with IAAFT the best calibrated overall. The full comparison across all six methods is reported in Fig S10. These findings establish IAAFT as the recommended default surrogate.

**Fig 3.**
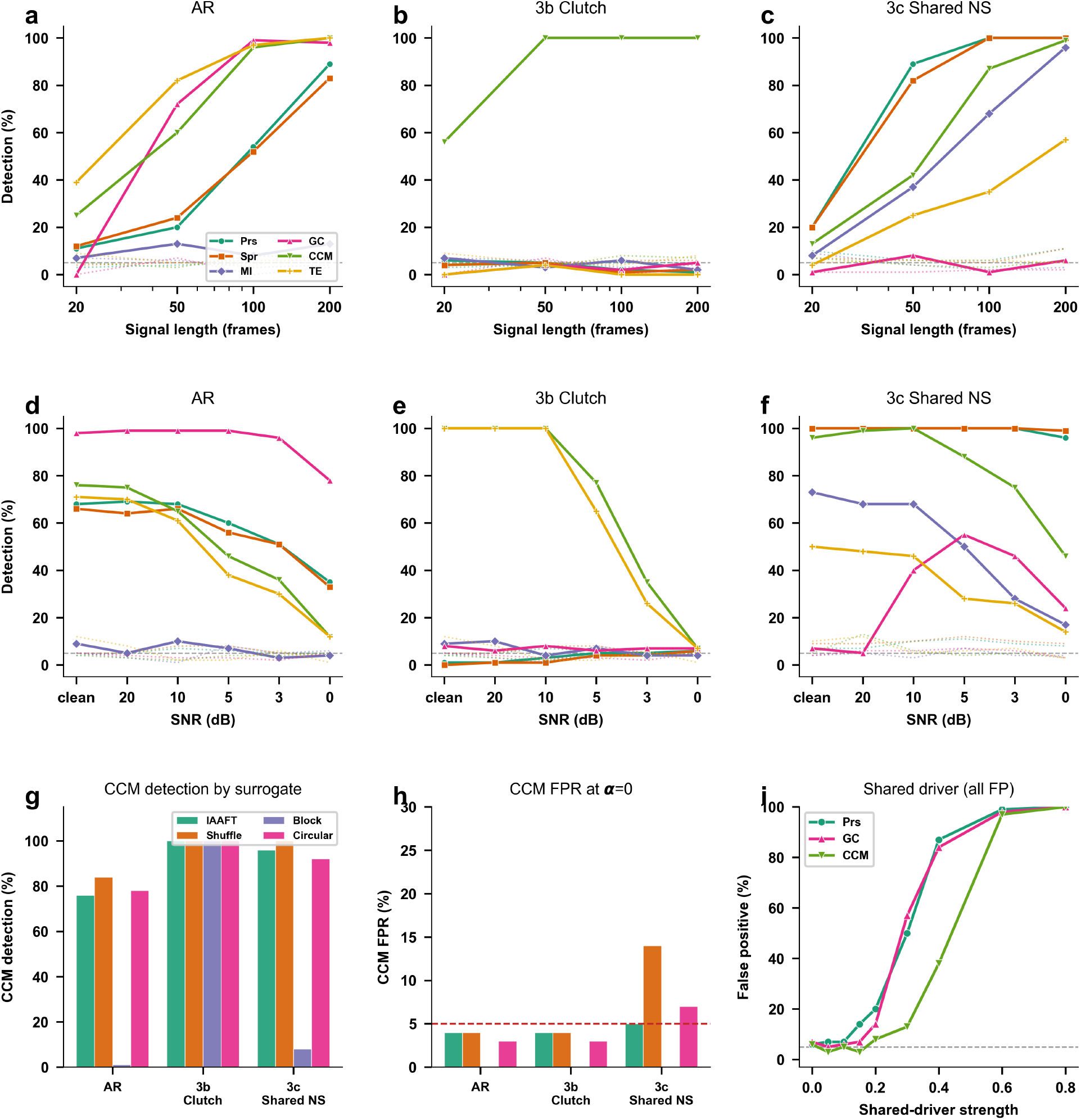
Robustness to signal length, observation noise, surrogate choice, and shared drivers. (**a–c**) Detection rate versus signal length (100 pairs, 1000 IAAFT surrogates; dashed line marks 5%; faint dotted curves show the false positive rate at *α* = 0) for AR (a), biphasic clutch (b), and shared non-stationarity (c). GC and CCM recover with length on AR; CCM detects the clutch from *n* = 50; correlation and CCM recover the shared-non-stationary coupling. (**d–f** ) Detection versus observation noise (SNR in dB) for the same three systems. GC is most noise-robust on AR; CCM and TE fall off sharply on the clutch (100% to near-zero by 0 dB); correlation stays noise-resistant on shared non-stationarity. (**g**) CCM detection by surrogate method at each system’s moderate coupling. Block bootstrap collapses CCM—it preserves the local temporal structure that manifold reconstruction exploits—while IAAFT, shuffle, and circular are comparable. (**h**) CCM false positive rate at *α* = 0 by surrogate method; shuffle inflates the FPR on the shared-non-stationary null, while IAAFT stays controlled. (**i**) Shared-driver false positives versus driving strength: a common latent source drives *X* and *Y* with no direct coupling, so all detections are false positives. Pearson and GC climb steeply (87%, 84% at strong driving); CCM is more conservative (38%). Surrogates cannot remove a genuine common cause.

##### (4) Signal length

Given the limitations due to photobleaching and stage drift, short time series are ubiquitous in live-cell imaging (30–100 frames). We swept four lengths (*n* = 20, 50, 100, 200) across three challenging systems (linear AR, *α* = 0.60; biphasic clutch, *α* = 1.20; and shared non-stationarity, *α* = 0.10), each at moderate coupling (Fig 3a–c). At *n* = 20, GC collapsed entirely (≤1% across all three systems), reflecting insufficient samples for VAR estimation. CCM retained detection at *n* = 20 for the clutch (38%) and remained ≥98% from *n* = 50, whereas the other methods stayed at chance at short lengths; only TE recovered at longer lengths (30% at *n* = 100, 99% at *n* = 200). For linear AR coupling, GC recovered fastest (72% at *n* = 50) and reached 99% by *n* = 100; CCM required *n* ≥ 100 for comparable sensitivity (96%). In the shared non-stationarity system, Pearson and Spearman detected the deterministic coupling at short lengths (42–46% at *n* = 50), while CCM lagged and GC remained blind across all lengths (≤4%), given removal of the coupling signal through stationarization. False-positive rates stayed near nominal, with mild inflation only on shared non-stationarity (≤11%). Overall, these results confirm that each method remains suited to its regime even as short signals make detection harder: GC for linear coupling, CCM for nonlinear and non-monotonic coupling, and correlation for shared non-stationarity.

##### (5) Observation noise

Adding white Gaussian noise (down to 0 dB SNR) exposed how each method fails as signals degrade (Fig 3d–f). GC excelled at linear AR coupling (*α* = 0.30), holding 78% detection at 0 dB, whereas CCM and TE fell to 12%, and Pearson and Spearman degraded gracefully to 33–35%. In the biphasic clutch (*α* = 1.20), CCM and TE held perfect detection down to 10 dB but collapsed to 7% once noise overwhelmed the attractor below 5 dB. The shared non-stationarity system inverted this fragility: Pearson and Spearman stayed at 96–99% even at 0 dB, because the large deterministic trends that carry the coupling tower over both the stochastic dynamics and the noise. GC behaved non-monotonically here, detection rising from 7% on clean signals to a peak of 55% at 5 dB before collapsing again, possibly because moderate noise disrupts the non-stationarity that stationarization would otherwise strip. False-positive rates stayed ≤13% throughout. Even under noise, then, the trichotomy holds: each method stays strongest in its own regime.

##### (6) Shared-driver confounding

So far we have tested situations in which two signals were directly related. However, coupling can also be spurious when a common upstream cause drives both. To probe this, we designed a shared-driver system in which a latent variable *Z* feeds both signals at different lags (*X* ← *Z*(*t*−1), *Y* ← *Z*(*t*−2)), with no direct *X* ↔ *Y* coupling, so every detection is by construction a false positive (Fig 3i). Using IAAFT surrogates, most methods had acceptable false-positive rates at low driving strength (≤7% at *α* = 0). As the driving strength increased, however, false positives rose steeply, reaching 87% (Pearson), 84% (GC), and 38% (CCM) at *α* = 0.4, and the methods could no longer distinguish direct coupling from a shared driver. CCM was the most resistant of the three, but no method was immune. Ruling out a common driver therefore requires experimental control and physical understanding beyond statistical analysis, testing the suspected driver directly for coupling against each downstream signal.

#### Summary of ground truth results

The ground-truth benchmarking (Table 3) substantiates the central claim: no single method is valid across all coupling regimes, and the three method classes (correlation, Granger causality, and convergent cross-mapping) partition the regimes between them. Granger causality owns the linear, stochastic regime and is the most noise-robust there. CCM, by contrast, recovers the two regimes where GC fails, nonlinear (clutch) and deterministic non-stationary coupling, and also survives severe signal shortening. Correlation, on the other hand, is broadly useful but offers no direction of influence and fails wherever the coupling is not monotonic. Throughout, IAAFT surrogate testing is what confirms each detection is genuine, making it a precondition, not an optional refinement.

**Table 3.** Summary of ground truth benchmarking. Detection rate (% significant at *p <* 0.05, 1000 IAAFT surrogates, 100 pairs per condition). Bold: best performer per condition.

| Condition | Prs | Spr | MI | GC | CCM | TE |
| --- | --- | --- | --- | --- | --- | --- |
| (1) AR $\alpha=0.4$ | 90 | 85 | 12 | **100** | 99 | 99 |
| (2) Logistic $\alpha=0.4$ | 35 | 33 | 63 | **100** | **100** | **100** |
| (3a) Separable nonstat $\alpha=0.4$ | 16 | 15 | 4 | **100** | 11 | 8 |
| (3b) Clutch $\alpha=0.4$ | 4 | 3 | 1 | 6 | **100** | **100** |
| (3c) Shared nonstat $\alpha=0.1$ | **100** | **100** | 73 | 7 | 96 | 50 |
| (3d) Time-varying $\alpha=0.4$ | 45 | 41 | 8 | **87** | 32 | 34 |
| AR short ($n=50$, $\alpha=0.6$) | 20 | 24 | 13 | 72 | 60 | **82** |
| Clutch short ($n=50$, $\alpha=1.2$) | 2 | 4 | 9 | 1 | **98** | 1 |
| AR noisy (0 dB, $\alpha=0.3$) | 35 | 33 | 4 | **78** | 12 | 12 |
| Clutch noisy (0 dB, $\alpha=1.2$) | 6 | 6 | 4 | 7 | 7 | 7 |
| Shared nonstat noisy (0 dB, $\alpha=0.1$) | 96 | **99** | 17 | 24 | 46 | 14 |
| *False positive rates ($\alpha = 0$):* |  |  |  |  |  |  |
| AR null ($n=300$) | 4 | 5 | 5 | 5 | 4 | 12 |
| Clutch null ($n=300$) | 4 | 5 | 5 | 5 | 4 | 12 |
| Shared nonstat null ($n=300$) | 7 | 9 | 4 | 4 | 5 | 10 |

These patterns motivate a trichotomy of Granger causality, Spearman, and CCM as the interpretive axes of Signal Associa, applied jointly under surrogate significance. Reading their agreement places a system along a spectrum from linear to non-monotonic coupling. At the linear end, GC (after stationarization) and Spearman agree, marking genuine linear coupling in the stochastic dynamics. One step to the left, only GC is significant while Spearman falls silent, signalling linear coupling masked by additive non-stationarity (3a). Toward the nonlinear middle, Spearman and CCM detect coupling that GC misses, indicating a nonlinear relationship or one carried by deterministic, non-stationary components (3b, 3c). At the non-monotonic extreme, CCM alone succeeds while Spearman fails. In other words, a non-significant GC excludes only linear stationary coupling, not other forms, so Spearman serves as a directionless middle player that also detects some nonlinear coupling. A negative Spearman, however, does not exclude coupling either, since non-monotonic mechanisms are visible only to CCM. Finally, no method is immune to shared-driver confounding, where a common upstream cause drives both signals; false positives are low at weak driving but rise steeply with driving strength, so ruling out a shared driver requires experimental control and physical reasoning, not statistics alone.

### Application 1: Natural killer cell collective coordination in tumor microenvironments

Natural killer (NK) cells are cytotoxic effectors of the innate immune system that patrol tissues in search of malignant and infected cells [20]. Recent imaging has shown that they do not act as isolated hunters: activated NK cells secrete diffusive chemokines that recruit distal NK cells to tumor targets, producing collective swarming through homotypic CCR5 signaling [21], and NK cells have been observed to accumulate cooperatively at specific tumor regions [22]. Despite these reports supporting that NK migration is coupled at the population level, no quantitative method has been established to assess it.

We applied Signal Associa to address this. NK cells were imaged in a micro-physiological tumor-vascular interface system [23] co-cultured with three colorectal cancer lines, the microsatellite-stable (MSS) HT29 and the microsatellite-instability-high (MSI-H; immunologically hot) LoVo and SW48, in 13 movies, one per chip, each chip from an independent experiment (HT29: 5, LoVo: 4, SW48: 4; 22.7 h at 5-min intervals; see Materials and methods). The analysis pipeline is illustrated on a representative HT29 movie in Fig 4: cells were segmented per frame (Fig 4a) and linked into trajectories with gap closing (Fig 4b). Trajectories of at least 20 frames were retained for analysis, 5,387 in total across the 13 movies (HT29: 1,829, LoVo: 2,353, SW48: 1,205).

**Fig 4.**
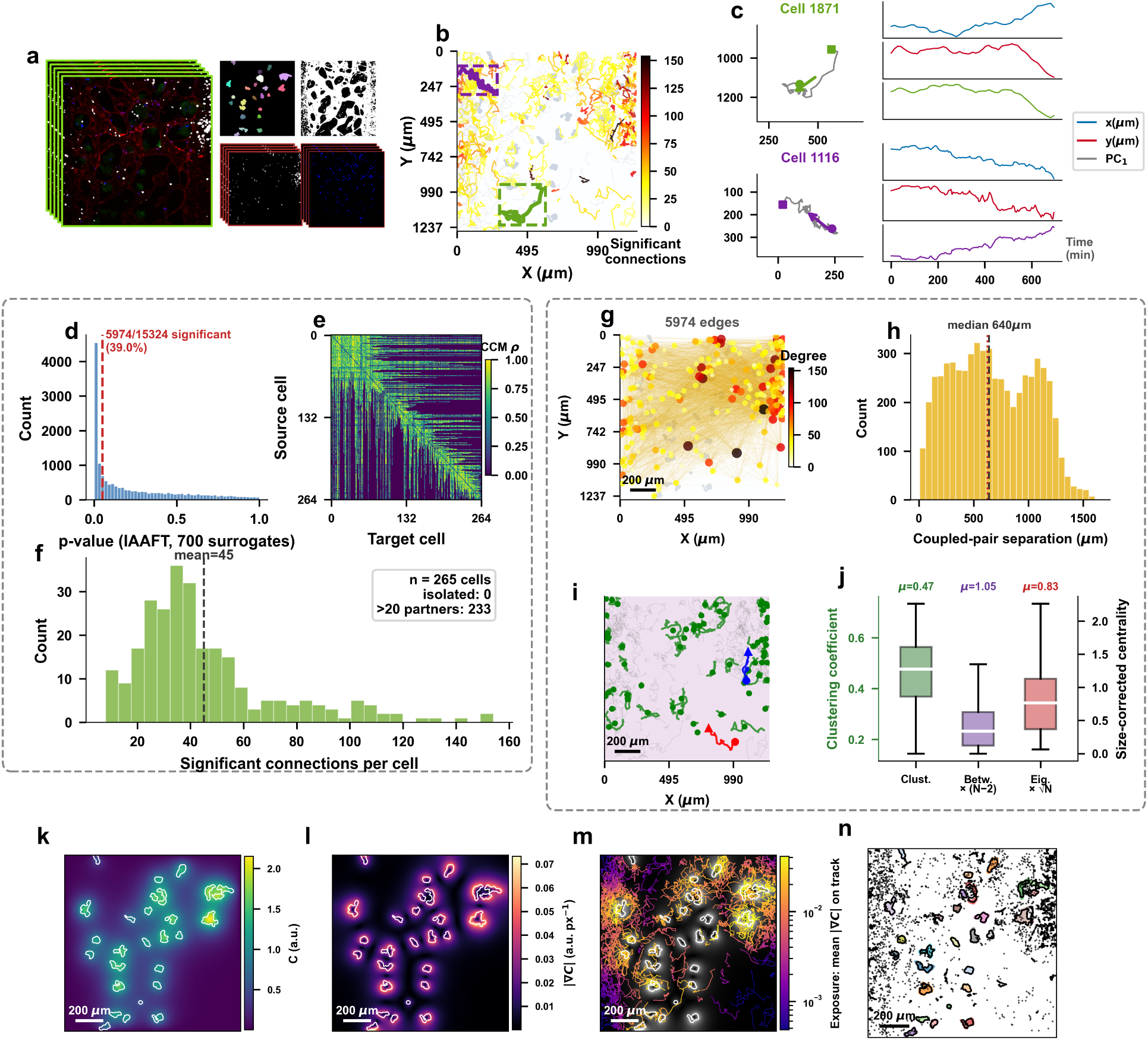
Application 1: NK coupling pipeline and network (representative HT29 movie) and tumor exposure. (**a**) Four-channel live-cell imaging of NK cells (white) with HT29 cancer cells (green) on endothelial monolayers (red), and segmentation masks. (**b**) NK cell trajectories colored by coupling degree. (**c**) PCA-based signal extraction: 2D trajectory projected onto PC_1_. (**d**) Distribution of CCM p-values (15,324 pairs, 700 IAAFT surrogates); 39% significant. (**e**) Influence matrix (265 × 265; color: cross-map *ρ*). (**f** ) Degree distribution. (**g**) Spatial coupling network (scale bar 200 *µ*m). (**h**) Physical separation of significantly coupled pairs; median 640 *µ*m. (**i**) Coupling at a distance: two cells 614 *µ*m apart that are not directly coupled, connected through intermediate cells coupled to both (green). (**j**) Distributions of the clustering coefficient (left axis) and of the size-corrected centralities, betweenness centrality ×(*N* − 2) and eigenvector centrality ×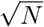(right axis), for the network of migrating cells (track extent ≥100 *µ*m) used in Fig 5; *µ*, means. (**k**) Steady-state Keller–Segel tumor field *C*(*x*), (*µ* − *D*_*C*_∇^2^)*C* = *S*_*tumor*_, with tumor outlines (white). (**l**) Gradient magnitude |∇*C*|, the local tumor signal a cell senses. (**m**) NK tracks colored by exposure, the mean |∇*C*| along each track (log scale). (**n**) Tumor regions (contours enlarged 1.3×; colors) and NK positions (black) used for the tumor-level analysis. Scale bars, 200 *µ*m.

If NK cells coordinate their motion as reported, that mutual influence should be visible in their migration dynamics. We therefore ask whether the trajectory of one NK cell can predict changes in the motion of another. To obtain one representative signal per trajectory, we compressed its *x*(*t*) and *y*(*t*) coordinates into a single time series using principal component analysis (PCA; Fig 4c). Repeating this test over all cell pairs with sufficient temporal overlap yields a network of influence whose structure reveals coordination at both local and global scales. We build this network for each of the 13 movies and assess how the tumor microenvironment reshapes its metrics.

Based on the ground-truth benchmarking, we selected CCM because NK trajectories are short (20–273 frames), nonlinear with cycles and linear trends, and deterministic, the regime where CCM is most appropriate. We nonetheless examine the pattern of agreement across the trichotomy to confirm this assignment.

#### Coupling network reveals dense collective coordination

Testing every NK-cell pair for coupling in the representative HT29 movie, a substantial fraction, 5,974 of 15,324 pairs (39.0%; pairs with sufficient temporal overlap, out of 34,980 in total), showed strong cross-map correlations (mean *ρ* = 0.883) that remained significant (*p <* 0.05) against 700 randomized surrogates (Fig 4d,e). This was not specific to one movie: the fraction of coupled pairs was 37–42% in every one of the 13 movies, in all three tumor types. Within the representative movie, no cell was isolated; every cell was coupled to at least one other (Fig 4f).

This motivates treating the population as a single influence network, with an edge between every pair whose coupling survives surrogate testing (Fig 4g). The node degree, the number of cells to which a given cell is significantly coupled, averaged 45 (17% of the population). Coupled cells were not necessarily neighbors: coupled pairs were as far apart as uncoupled ones (median 640 versus 639 *µ*m; Mann-Whitney *p* = 0.97), spanning the entire field with 51% of coupled pairs more than half a field width apart (Fig 4h), and the same held across the 13 movies (coupled pairs a median 1.6% closer than uncoupled pairs). Such coupling at a distance can also be relayed through the network: two cells 614 *µ*m apart that were not directly coupled were connected through intermediate cells coupled to both (Fig 4i).

#### Coordination is universal; its integration at matched tumor signal is tumor-dependent

The coupling network allows each NK cell to be characterized by its role in the collective. We use three complementary graph metrics that span spatial scales. The clustering coefficient measures how tightly a cell’s coupling partners are coupled among themselves, a *local* property [24]. This local grouping was not a by-product of network density: in all 13 movies, clustering exceeded that of randomized networks with the same degree of every cell (1.07–1.47-fold; median 1.20) and of random networks with the same density (1.30–2.39-fold; Wilcoxon *p* = 1.2 × 10^*−*4^ for both). Betweenness centrality measures how often a cell lies on the shortest coupling paths between other cells, i.e., its role as a *relay* between distant cells (as in Fig 4i). Eigenvector centrality scores a cell high if it is coupled to cells that are themselves highly connected [25]; because the definition is recursive, it reflects a cell’s influence across the whole network, a *global* property. Both centralities depend on network size: unit-norm eigenvector centrality decreases as 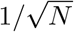, and the network mean of normalized betweenness equals (*L* − 1)*/*(*N* − 2), where *L* is the mean shortest-path length. We therefore multiplied eigenvector centrality by 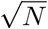 and betweenness by (*N* − 2) so that networks of different sizes (148–689 cells) can be compared. Fig 4j shows the three distributions for the representative HT29 movie.

Across all 13 movies, the coordination rate was nearly constant (37–42% of tested pairs), as was coupling strength (mean *ρ* of coupled pairs 0.88–0.91 per movie), so collective coordination itself is a conserved feature of NK migration in these systems. NK cells, however, experience very different amounts of tumor-derived signal depending on tumor size and proximity. We modeled this signal as the steady-state Keller–Segel field generated by the tumors (Fig 4k) and defined each cell’s exposure as the mean gradient magnitude |∇*C*| along its track (Fig 4l,m). Exposure was highest in the MSS line HT29 (median 0.028 versus 0.020 and 0.019 in LoVo and SW48), whose fields contain more tumor. To separate the effect of tumor type from that of exposure, we compared cells at matched exposure: all cells were placed in common exposure bins and the three metrics were compared between tumor types within each bin, with movies as the units of replication (Fig 5a–c). In every tumor type the two centralities increased with exposure (Spearman *ρ* over cells +0.07 to +0.11, all *p <* 0.05), whereas clustering did not. At any given exposure, cells around MSI-H tumors were more globally integrated than HT29 cells: for eigenvector centrality × 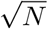, the MSI-H mean was higher in all four bins (MSI-H minus HT29 = +0.059, exact permutation *p* = 0.008 over the 13 movies), betweenness ×(*N* − 2) tended to be higher (+0.082, *p* = 0.05), and clustering differed little (+0.016, *p* = 0.07). Networks were built from migrating cells (track extent ≥100 *µ*m; 84% of trajectories); the eigenvector centrality difference persisted when all cells were included (+0.045, *p* = 0.03). Thus HT29 cells experienced the strongest tumor field, yet at the same exposure they were less integrated into the coupling network than cells around MSI-H tumors.

**Fig 5.**
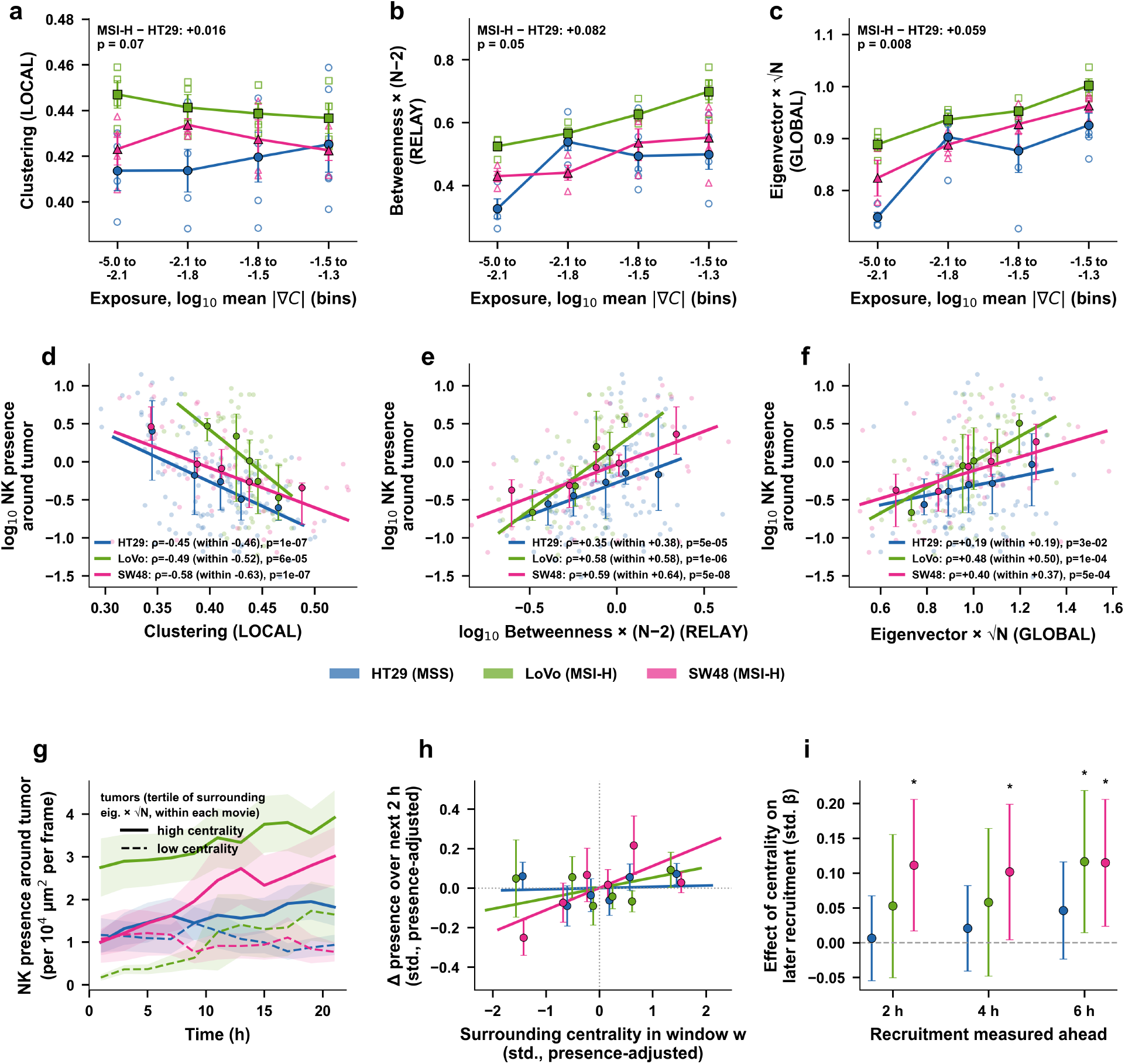
Application 1: NK network coordination at matched tumor exposure, and its link to NK recruitment. (13 movies; HT29 (MSS): 5, LoVo and SW48 (MSI-H): 4 each; networks built from migrating cells, track extent ≥100 *µ*m). (**a–c**) Network metrics versus each cell’s own tumor exposure, the mean Keller–Segel gradient |∇*C*| along its track: clustering coefficient (local, a), betweenness centrality ×(*N* − 2) (relay, b) and eigenvector centrality ×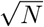 (global, c). Cells are placed in common quantile bins of log_10_ exposure (bin ranges on the axis; the sparsely populated top bin is omitted). Open markers, mean per movie (cells beyond 1.5 IQR within type and bin excluded); filled markers and lines, mean ± s.e.m. over movies. Inset: MSI-H minus HT29 averaged over bins, with the exact permutation *p* over all assignments of the 13 movies to the two groups. (**d–f** ) Per tumor (262 tumors; region = tumor contour enlarged 1.3×): NK presence (log_10_) versus the time-weighted metric of the NK cells around the tumor; lines are Theil–Sen fits, points with bars are medians and interquartile ranges per quintile of the metric, *ρ* is Spearman’s correlation over all tumors of a type and, in parentheses, averaged within movies (Fisher *z*). (**g**) NK presence around tumors over time (2-h windows; mean ± s.e.m. over tumors) for tumors in the top (solid) and bottom (dashed) tertile of surrounding eigenvector centrality ×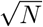, tertiles assigned within each movie. (**h**) The data behind panel i, for the 2-h horizon: change in a tumor’s NK presence over the next 2 h versus the eigenvector centrality ×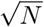 of its surrounding NK cells in the current window, both after removing what the current presence explains (partial residuals; variables centred within movie and standardized). Points, mean ± s.e.m. per sextile of centrality over tumor-windows; lines, the fitted slopes, which are the *β* values shown in i. (**i**) Does integration predict recruitment? Standardized effect of the surrounding eigenvector centrality ×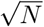 in one window on the change in NK presence over the next 2, 4 or 6 h, adjusted for the current presence (linear mixed model with a tumor random intercept; all variables centred within movie); points, *β* ± 95% CI; ^*\**^*p <* 0.05.

We next asked whether this coordination is linked to NK recruitment around individual tumors. Each tumor contour was enlarged by a factor of 1.3 about its centroid (Fig 4n), and for every tumor we measured NK presence (NK cell-frames per unit area per frame) and the network metrics of the NK cells around it, weighted by the time each cell spent there (262 tumors; HT29: 130, LoVo: 60, SW48: 72). Around tumors that recruited more NK cells, local clustering was lower in all three tumor types (Spearman *ρ* = −0.45, −0.49, −0.58 for HT29, LoVo, SW48; all *p <* 10^*−*4^; Fig 5d), whereas the relay and global roles were higher (betweenness ×(*N* − 2): *ρ* = 0.35, 0.58, 0.59; eigenvector centrality × 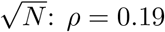, 0.48, 0.40; Fig 5e,f). These correlations were as strong within individual movies as across them (Fig 5d–f, values in parentheses), so they do not reflect differences in NK numbers between movies. As recruitment rises, NK networks thus shift from tight local groups toward relay and global integration, and this link tended to be stronger around MSI-H tumors than around HT29 tumors (eigenvector centrality: HT29 versus MSI-H, Fisher *z, p* = 0.03).

The tumor-level correlations do not show which comes first: does an integrated NK network attract more NK cells, or does a crowded tumor make its NK cells more integrated? To answer this, we followed each tumor over time. At the start of the recordings, NK presence around SW48 and HT29 tumors was about one cell per 10^4^ *µ*m^2^ per frame, whether their surrounding network was highly integrated or not (tumor regions are typically about 10^4^ *µ*m^2^, so this is roughly one NK cell present at a time). Over the next 22 h, tumors with a highly integrated network (top third within each movie) rose to 3.0 (SW48) and 1.8 (HT29) cells per 10^4^ *µ*m^2^, while tumors with a poorly integrated network stayed below 1 (Fig 5g). LoVo tumors showed the same pattern at a higher level, with integrated tumors rising from 2.8 to 3.9 cells per 10^4^ *µ*m^2^. Because the tertiles are defined by centrality over the whole movie, these curves are descriptive; the temporal order is tested by the model in Fig 5h,i. In total, 28 different NK cells visited a typical integrated LoVo tumor during the recording, compared with 10 for a poorly integrated one. We then compared tumors that held the same number of NK cells at a given time. Those with a more integrated network gained more NK cells over the next hours around SW48 tumors (at 2, 4 and 6 h; standardized *β* = 0.10–0.12, *p* = 0.01–0.04) and around LoVo tumors (at 6 h; *β* = 0.12, *p* = 0.03), but not around HT29 tumors (*β* ≤ 0.05; Fig 5h,i). Around MSI-H tumors, a more integrated NK network therefore tends to precede, modestly but consistently, further NK recruitment.

#### Multi-method comparison and regime classification

To test how the trichotomy plays out on real data, we applied all five method families to every one of the 15,324 HT29 pairs: classic Pearson and Spearman (lag 0), lag-scanned Pearson (Danuser), Granger causality (auto-stationarized), and CCM, each with IAAFT surrogate significance. As anticipated from the ground truth systems, the lag-scanned Pearson method generously flagged 94.5% of all pairs as coupled at its parametric threshold, but only 32.7% survived surrogate testing, underscoring the necessity of surrogate control. Granger causality, by contrast, identified almost no coupling (6.6%), and its detections barely overlapped those of any correlation method. This near-absence of GC-detectable coupling indicates that little of the association resides in the stationary stochastic dynamics that GC models; after stationarization removed the trends and cycles, GC had little left to detect.

CCM, on the other hand, detected 39.0% of pairs (Fig 6), about half of which (54.0%) were also recovered by lag-scanned Pearson, consistent with coupling carried by shared cycles and trends in the ground truth systems; more importantly, the remaining 46.0% were detected by CCM but not by any correlation method, representing nonlinear or non-monotonic associations invisible to linear approaches. A smaller set of pairs (11.6%) was detected by lag-scanned Pearson but not CCM, consistent with weak, largely linear couplings on short trajectories where CCM struggles. The overall picture, dominated by nonlinear and deterministic coupling with almost no purely linear stochastic component, is the regime signature that motivated selecting CCM for this system, and the multi-method analysis by Signal Associa provided further insight into it.

**Fig 6.**
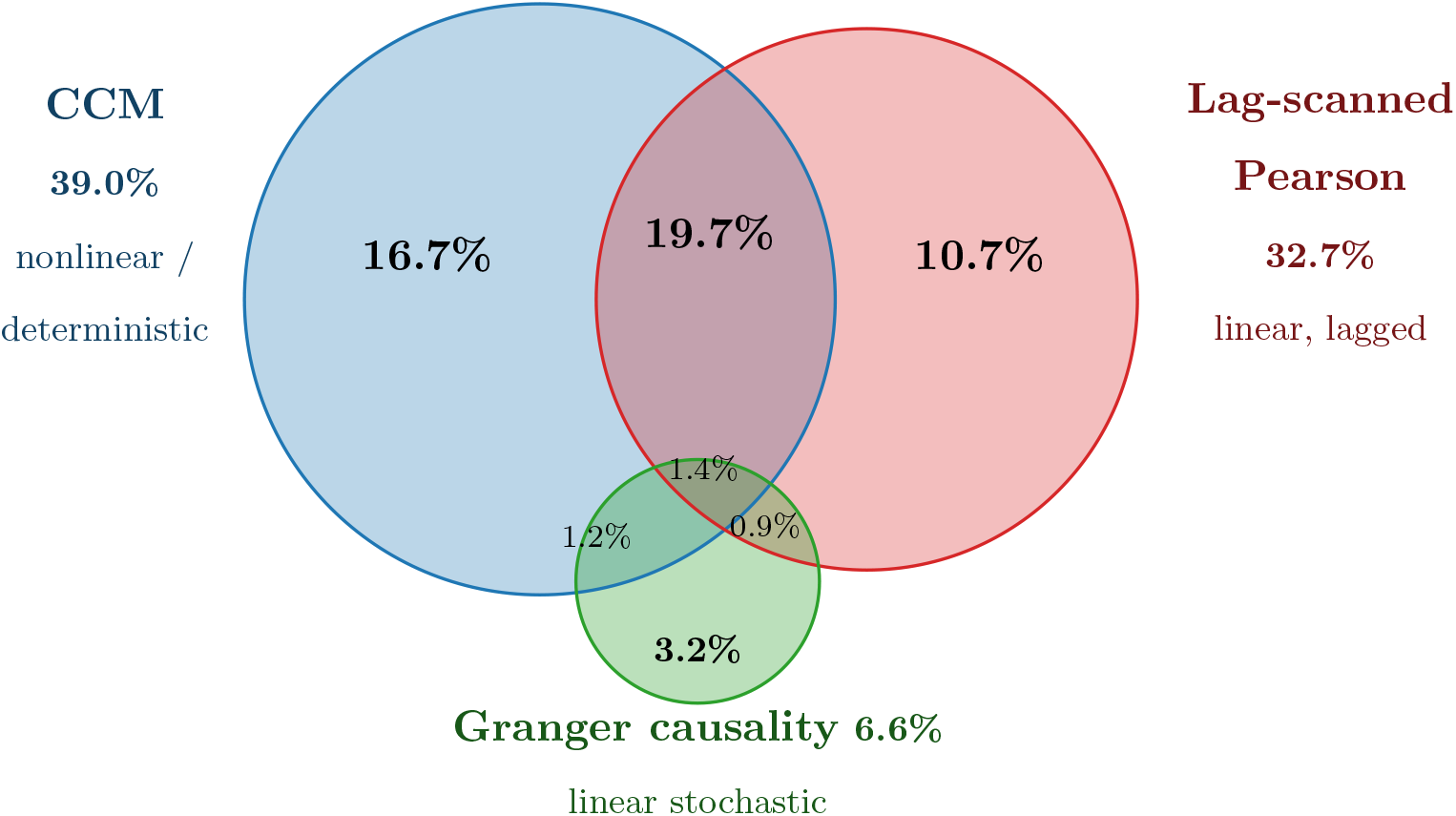
Multi-method detection overlap on NK HT29 pairs. (15,324 pairs, IAAFT surrogate significance). Circle area is proportional to each method’s detection rate; overlap regions give the percentage of all pairs. CCM (39.0%) captures the largest set, including a nonlinear/deterministic component invisible to the other methods, while Granger causality (6.6%) is small and nearly disjoint. The remaining 46.2% of pairs showed no coupling by any method.

### Application 2: Vinculin–FAK phosphorylation coupling varies with stiffness and cell region

Focal adhesions transmit the mechanical state of the extracellular matrix to intracellular signaling, with focal adhesion kinase (FAK) as a central signaling node [26]. However, there have been contradictory reports about the interplay of FAK, talin, and stiffness. One line of evidence places FAK on a separate axis from the mechanosensor talin: FAK and Src are activated independently of adhesion complex composition [27]. On the other hand, FAK and talin recruit one another at nascent adhesions [28] upon talin unfolding induced by force, and the linear relationship between FAK Y397 phosphorylation and traction at individual adhesions is abolished once the E1015A mutation disrupts FAK-talin binding [29]. While FAK Y397 phosphorylation has been reported to increase with higher stiffness [26], the classical model predicts that adhesion turnover requires increased FAK phosphorylation to suppress RhoA [30]; on stiffer substrates, however, transmitting more force requires higher RhoA activity, which contradicts the concurrent increase of FAK phosphorylation and RhoA on stiff substrates [31].

In contrast to perturbation studies, which eliminate spatial and temporal information, we addressed the first part of this contradiction by imaging live cells expressing a FRET-based FAK biosensor (Fig 7a), whose EGFP/FRET ratio reports FAK Y397 phosphorylation [26], together with fluorescently tagged vinculin (Fig 7a), a marker of talin unfolding under tension, to investigate their dynamic interaction and possible mutual influence. After segmenting individual focal adhesions and tracking them over time, we read both signals along each trajectory as paired time series (Fig 7a– c). We then applied Signal Associa to detect coupling between vinculin recruitment and FAK activation at single adhesions on soft (0.6 kPa) and stiff (6 kPa) substrates to account for varying force-generation capacity.

**Fig 7.**
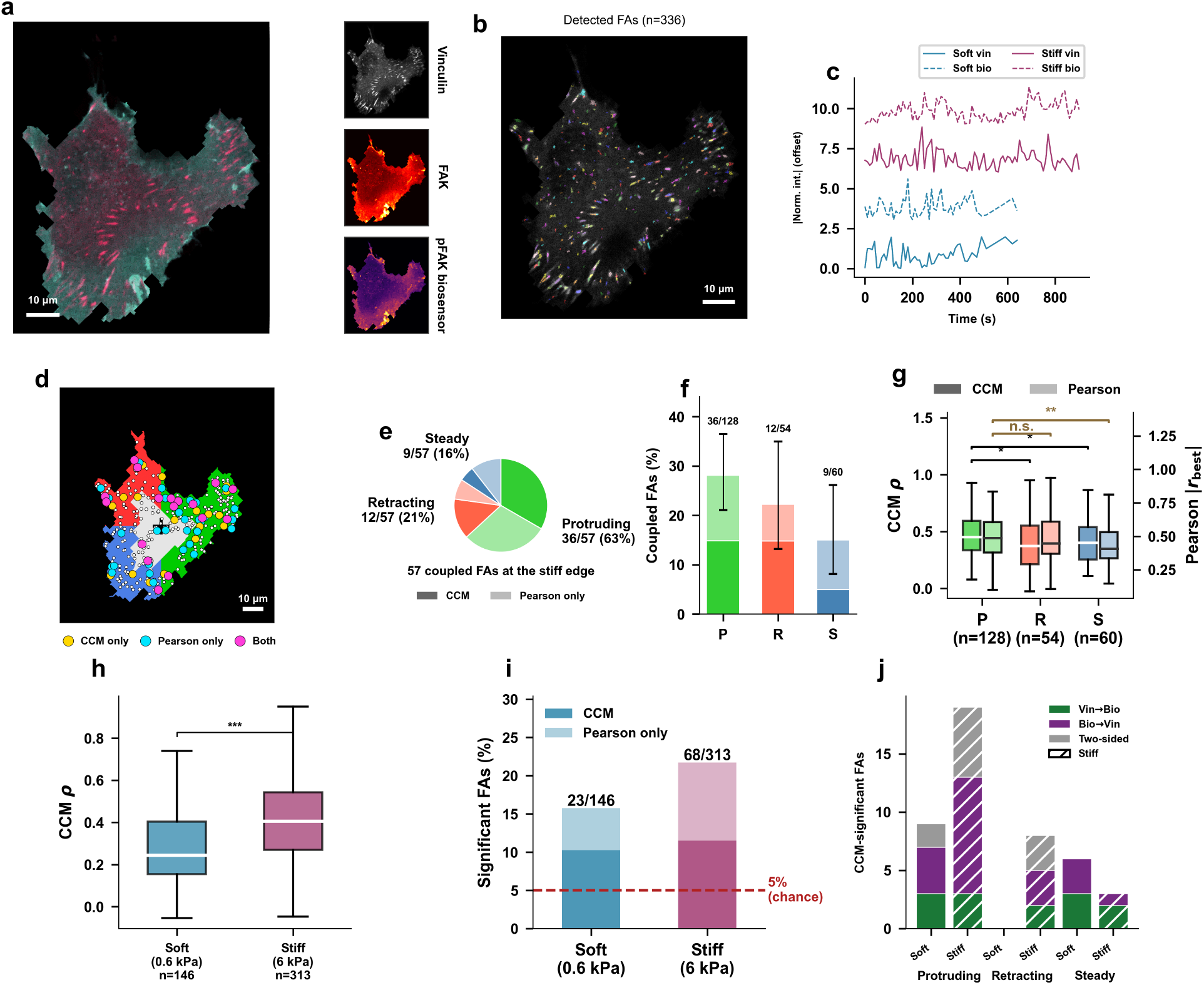
Application 2: ECM stiffness-dependent vinculin–FAK phosphorylation coupling at individual focal adhesions. (**a**) Multi-channel imaging on stiff substrate (6 kPa): composite, vinculin, FAK, and pFAK biosensor channels. (**b**) Focal adhesions detected in a single frame (*n* = 336), colored by identity. (**c**) Representative time series on soft (blue) and stiff (purple). (**d**) Cell edge regions (a ∼12 *µ*m band; the gray interior is excluded from panels d–g): protruding (green), retracting (red), steady (blue). Dots are tested FAs (tracks long enough for CCM); significantly coupled FAs are marked by method: CCM only (gold), Pearson only (cyan), both (magenta). (**e**) Significantly coupled edge FAs by region (union of CCM and lag-scanned Pearson; *n* = 57), each wedge split into a CCM-significant (darker) and Pearson-only (lighter) share; inner labels give each region’s share of coupled FAs (e.g., 36/57), outer labels the fraction of tested FAs in that region that were coupled (e.g., 36/128). (**f** ) Fraction of tested FAs coupled per region (CCM, darker; Pearson-only, lighter), with Wilson 95% confidence intervals. (**g**) Regional coupling strength (stiff; tested FAs): CCM *ρ* (left axis) and Pearson |*r*_best_| (right axis). Protruding *>* retracting and *>* steady for CCM (one-sided Mann-Whitney *p* = 0.012 and 0.025); for Pearson, protruding *>* steady (*p* = 0.006), protruding versus retracting n.s. (**h**) Overall CCM *ρ* of tested FAs: stiff *>* soft (*p <* 0.0001). (**i**) Detection rate, stacked as CCM (base) and Pearson-only (top): soft 15.8% (23/146), stiff 21.7% (68/313 tested FAs over the whole cell, including interior FAs); dashed line, 5% chance. (**j**) Directionality of CCM-significant FAs by region and stiffness: number of adhesions whose cross-map skill is stronger in one direction (difference *>* 0.10; Vin→Bio or Bio→Vin) or comparable in both (two-sided); hatched bars, stiff.

Applying the trichotomy to the focal-adhesion data placed this system in a different regime than the NK population. As before, the lag-scanned Pearson method flagged a large fraction of adhesions at its parametric threshold (49% soft, 71% stiff), but after surrogate testing only a small fraction of adhesions was coupled by either method (15.8% soft, 21.7% stiff; Fig 7i), and Granger causality detected almost nothing in either condition (below 4%), indicating that little of the coupling resides in stationary stochastic dynamics. Unlike the NK trajectories, however, CCM and the correlation-based method recovered overlapping sets of adhesions: of the coupled adhesions on stiff substrates, 17 were detected by both, against 19 by CCM alone and 32 by Pearson alone. This agreement of CCM and correlation while Granger causality stays silent indicates that vinculin and pFAK co-vary largely monotonically through shared cycles and trends, placing focal-adhesion coupling in the linear, lagged corner of the trichotomy, opposite the nonlinear, deterministic corner occupied by the NK population.

The coupling that did occur was not uniform across the cell. We divided the cell edge into protruding, retracting and steady regions (a ∼12 *µ*m band; adhesions in the cell interior were excluded) and mapped the significantly coupled adhesions, which CCM and Pearson flagged at largely the same locations (Fig 7d). Of the 57 coupled adhesions at the edge of the cell on the stiff substrate, 36 (63%) were at protrusions (Fig 7e). Protrusions also had the highest fraction of coupled adhesions: 28% (36/128 tested adhesions), versus 22% (12/54) at retracting and 15% (9/60) at steady edges, although not significantly (*χ*^2^ *p* = 0.14; Fig 7f). Coupling was also strongest at protrusions (median CCM *ρ* = 0.451 versus 0.372 retracting and 0.400 steady; one-sided Mann-Whitney *p* = 0.012 and 0.025), and Pearson coupling agreed for protruding versus steady (*p* = 0.006) but not versus retracting (*p* = 0.20; Fig 7g). Protruding edges are sites of nascent adhesion and rapid force development, and coupling was strongest exactly there.

Stiffness strengthened coupling rather than making it more frequent. The median CCM *ρ* rose from 0.245 on soft to 0.406 on stiff substrates (*p <* 0.0001; Fig 7h), whereas the fraction of coupled adhesions was similar (15.8% versus 21.7%, Fisher *p* = 0.17; CCM alone 10.3% versus 11.5%; Fig 7i). At the level of individual adhesions, at protruding edges on stiff substrates FAK phosphorylation tended to lead vinculin (10 of 19 significant adhesions Bio→Vin versus 3 Vin→Bio; binomial *p* = 0.09). Steady and retracting edges had too few coupled adhesions (3 and 8) for a dominant direction to emerge (Fig 7j).

Returning to the contradiction that motivated this analysis, our per-adhesion measurements and coupling assessment show that vinculin and FAK phosphorylation are not coupled by a single fixed relationship, but vary by location and stiffness. Coupling was weak on soft substrates and became stronger, though not more frequent, on stiff substrates. More strikingly, within the stiff condition it was strongest at protruding edges, where coupled adhesions also tended to be most common, although not significantly so. This stiffness- and location-dependence is consistent with the view that FAK activation depends on talin mechanosensitivity only under a certain mechanical condition [29], and it reconciles the two opposing lines of evidence: the apparent independence of FAK from talin [27] and their interdependence [28] may reflect adhesions sampled in different mechanical regimes rather than a single fixed pathway.

By recovering this structure from observational time series alone, Signal Associa provides a way to test where and when the structural and signaling arms of the adhesion are linked, a question central to how cells sense substrate stiffness.

## Discussion

Our ground truth benchmarking shows that the coupling landscape partitions into three regimes, each with a clear champion, and the six ground truth systems map cleanly onto this partition. Correlation (Pearson, Spearman), while unable to provide information about the direction of influence, was the most sensitive for monotonic, stationary coupling: it detected the linear AR system (1) at high coupling and the shared deterministic system (3c) at 100%. Nonetheless, it failed to detect non-monotonic clutch coupling (3b, ≤4%), where the mean is not a good representative of the dynamics. In the same way, Granger causality dominated linear stochastic dynamics: it reached 100% on AR (1) and, after stationarization, on separable non-stationarity (3a), and 87% on time-varying coupling (3d); but it was blind when the coupling lived in the non-stationary components of the signal that stationarization removes (3c, ≤8%). Convergent cross-mapping completed the trichotomy: it detected both regimes that defeat Granger causality, the clutch (3b, 100%) and shared deterministic coupling (3c, 96%), being the only method to recover the clutch across the full coupling range, and it was the most robust to short signals. The shared-driver sensitivity experiment exposed a common limit of all methods: when a hidden variable drives two downstream signals at once, they cannot distinguish direct coupling between the two signals from being driven by a common influence. CCM was the most sturdy against this artifact. Given this trichotomy, the choice of method should follow the expected coupling regime rather than convention.

This trichotomy carries a practical consequence: coupling methods cannot be applied blindly. Choosing the wrong method for a given regime does not merely reduce sensitivity; it produces systematically misleading results. Correlation applied to non-monotonic coupling returns values near zero regardless of coupling strength, not because coupling is absent but because positive and negative contributions cancel. Granger causality applied to deterministic coupling returns null results after stationarization, not because the signals are independent but because the coupling was carried by the very components that were removed. The appropriate method therefore depends on the expected coupling regime: whether the coupling is linear or nonlinear, monotonic or non-monotonic, and whether it resides in stochastic fluctuations or in shared deterministic structure such as trends and oscillations. Because no single method spans these regimes, running them together and reading their agreement, rather than committing to one in advance, is what makes the regime identifiable from the data.

The largest false-positive occurrence we observed came from the lag-scanned Pearson cross-correlation widely used in live-cell biosensor and protrusion imaging: at its parametric 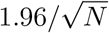 threshold, false positive rates reached 72–92% on uncoupled autocorrelated signals, because scanning over lags and taking the peak compounds the autocorrelation that already violates the test’s independence assumption. The classical single-lag methods were much less affected. Granger causality’s parametric test remained well controlled (≤9%), and Pearson and Spearman stayed near nominal on stationary systems (≤6%), rising only modestly, to about 11%, on the non-stationary systems (3a, 3c) where trends and cycles inflate the raw correlation. IAAFT surrogate testing, which randomizes the signals in a way that preserves each signal’s power spectrum and amplitude distribution while destroying cross-signal coupling, brought these rates down to about 5–10% across methods and systems. Given these results, surrogate-based null testing, specifically IAAFT, should be preferred over classical distribution-based tests for genuine coupling detection and accurate biological findings.

The two applications show the trichotomy playing out on real data, with the two systems landing in opposite corners. In NK-cell migration, lag-scanned Pearson flagged nearly all pairs as coupled (94.5%), but surrogate testing kept only a third (32.7%) and Granger causality detected almost none (6.6%), while CCM recovered the largest set (39.0%), nearly half of it invisible to any correlation method. This pattern, CCM dominant with partial overlap with correlation whereas GC is silent, places NK coordination in the nonlinear, non-monotonic region of the spectrum. On the other hand, focal-adhesion mechanotransduction landed in the opposite corner: correlation and CCM agreed on overlapping sets of adhesions while Granger causality again stayed silent, the signature of coupling carried monotonically through shared cycles and trends, which are removed by stationarization. In both systems, reading the pattern of agreement across methods helped identify the regime, and showed that conducting all methods simultaneously is necessary.

Beyond method selection, coupling analysis gave insight into biological behavior that no single measurement captures. In the first application, NK cells formed dense coupling networks at the same rate in every movie and tumor type, so coordination itself appears to be a conserved feature of NK migration. What differed between tumors was how the network was organized: at the same exposure to tumor-derived signal, NK cells around MSI-H tumors were more globally integrated than around MSS tumors. Around individual tumors, higher NK recruitment went together with fewer tight local groups and more cells acting as relays and global hubs, and this organization was not merely a consequence of crowding: around MSI-H tumors, a more integrated surrounding network tended to precede the arrival of further NK cells. Because only one MSS line was studied, an HT29-specific effect cannot be excluded, and additional MSS lines will be needed to attribute these differences to microsatellite status. In the second application, combining coupling analysis through Signal Associa with spatial mapping across cell regions clarified the apparently contradictory role of FAK in mechanosensing: vinculin–FAK coupling was not fixed but depended on mechanical context: it was strengthened on stiff substrates and strongest at protruding edges, reconciling the conflicting reports of FAK-talin independence and interdependence as adhesions sampled in different mechanical regimes.

These results point to a few practical considerations. The methods are best run together, with IAAFT surrogation, so that they complement each other and the results can be read in a more insightful and informative way. Agreement between correlation and Granger causality is consistent with linear stochastic coupling; coordination of CCM and Pearson points to coupling residing in cycles and trends; and detection by CCM alone indicates non-monotonic dynamics. Disagreement, therefore, helps localize the regime rather than signaling an error. Real recordings, however, are short and noisy, and both interact with method choice. Under observation noise, GC was the most robust of the methods (retaining 78% detection at 0 dB on the AR system), whereas CCM degraded at high noise levels. Signal length pulls in the opposite direction: CCM recovered coupling from as few as 50 frames on nonlinear systems, while GC needs longer series to fit its autoregressive model. In practice the choice is rarely free: a short, clean nonlinear recording favors CCM, while a longer, noisier linear one favors GC, and where the regime is genuinely uncertain the agreement structure across methods is the most reliable guide.

The current framework has clear boundaries. Signal Associa operates on pairs of signals, and pairwise methods cannot distinguish direct coupling from indirect paths through a third variable, as the shared driver results (Pearson 87%, Spearman 82%, GC 84%, CCM 38% false positives) illustrate. In the NK application, some of the detected coupling may reflect a shared response to the same tumor cues rather than direct NK–NK signaling. Extending the framework to multivariate methods, including partial correlation, conditional transfer entropy, and multivariate CCM, would enable disentangling direct from mediated coupling in systems with more than two interacting components. Signal length also constrains performance: all methods require sufficient time frames and the true positive detection rate degrades below *n* = 50 even on well-characterized nonlinear systems. Importantly, short signals do not inflate false-positive rates; they remain reliable for ruling out coupling, but may fail to detect genuine associations at weak coupling strengths. Finally, GPU-accelerated surrogate testing requires NVIDIA hardware and CuPy, though CPU-only execution remains available at reduced throughput.

## Materials and methods

### Ground truth systems

To systematically evaluate detection methods across the coupling regimes encountered in biological time series—linear, nonlinear, confounded, and non-stationary—we designed six ground truth systems, each with known coupling structure and adjustable coupling strength *α*. Each system is motivated by a specific biological scenario and targets a known failure mode of one or more detection methods (see Coupling detection methods and Validation framework below for the methods and surrogate testing procedures used to analyze them).

#### (1) Coupled autoregressive process (linear)

Two linearly coupled stochastic signals with unidirectional coupling *X* → *Y* :

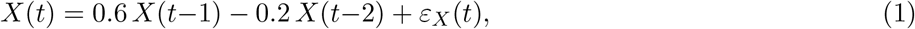

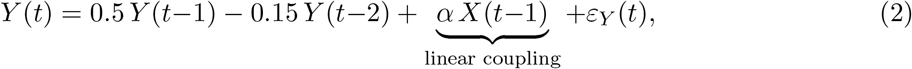

where *ε*_*X*_, *ε*_*Y*_ ∼ *N* (0, *σ*^2^=0.09) are independent Gaussian noise terms (*σ* = 0.3). The system is initialized with 100 burn-in steps that are discarded. This system is stationary and linear: Granger causality is expected to perform well, while CCM may show reduced sensitivity or incorrect directionality [32].

#### (2) Coupled logistic maps (nonlinear)

Following Sugihara et al. [3] (Eq. 1), the system uses bidirectional coupling:

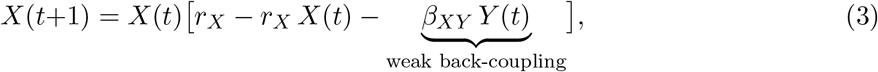

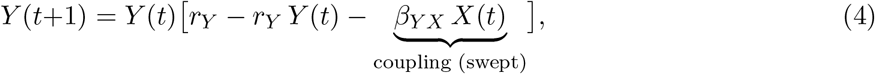

with *r*_*X*_ = 3.8, *r*_*Y*_ = 3.5 (both chaotic), *β*_*XY*_ = 0.02 (fixed weak back-coupling), and *β*_*Y X*_ swept as the coupling parameter *α*. Initial conditions are *X*(0) = 0.4+*δ, Y* (0) = 0.4+*δ* where *δ* ∼ *N* (0, 0.01). Values are clipped to [0.001, 0.999] to prevent divergence. The system uses 500 burn-in steps and *n* = 100 (realistic biological signal length). The bidirectional coupling produces state-dependent “mirage correlations”—the long-term Pearson correlation is near zero (*r* ≈ 0.04 at *β*_*Y X*_ = 0.1, *n* = 100) despite genuine dynamical coupling, because the chaotic dynamics produce alternating windows of positive, negative, and zero correlation [3].

#### (3) Non-stationarity and nonlinearity

Biological signals are rarely stationary, and the coupling between them may be nonlinear. These two challenges require different analytical strategies. We test both.

#### (3a) Separable non-coupled non-stationarity

The underlying coupling is the same linear AR mechanism as system 1 (Eqs. 1–2), but *X* is observed through additive non-stationary contamination:

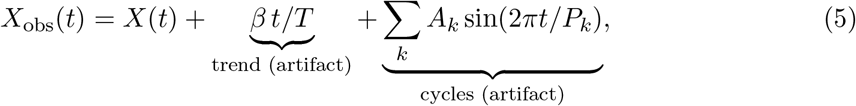

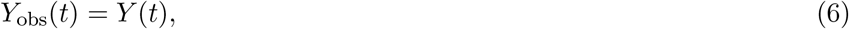

where *β* controls trend strength, *A*_*k*_ the cycle amplitudes, and *P*_*k*_ the cycle periods. Critically, *Y* couples to the *clean X*(*t*), not to *X*_obs_(*t*): the non-stationarity is independent of the coupling mechanism. The Stationarizer can therefore decompose and remove the trend and cycles (Eq. 29), recovering the stationary residual on which Granger causality operates correctly. This system demonstrates the Stationarizer’s value: without preprocessing, GC produces spurious results; with it, GC recovers the true coupling.

#### (3b) Biphasic clutch coupling (non-monotonic)

In the molecular clutch model of mechanotransduction [7], the coupling between actin retrograde flow and traction force is biphasic: at low flow speeds, molecular clutches engage and force builds proportionally; past a threshold, clutches rupture and force drops even as flow increases. Craig et al. [33] derived an analytical steady-state approximation for the ensemble-averaged traction:

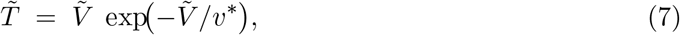

where 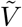 is the actin flow speed and *v*^*\**^ a characteristic velocity set by clutch stiffness and Bell-model parameters [34] (their Eq. 9 with *b* = 0, no myosin feedback). The linear factor 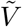 arises because clutch force grows as *F* = *κ*_*c*_ *v τ* (Hookean loading, Chan & Odde Eq. 2 [7]), and the exponential decay 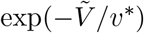 reflects Bell-law bond failure: 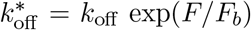 (Chan & Odde Eq. 1 [7]; Bell [34]), which shortens the mean clutch lifetime exponentially with flow speed.

Following Eq. (7), we define the coupling function as:

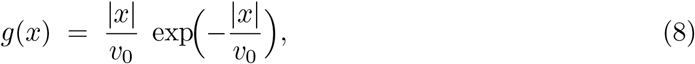

where |*x*| maps signed velocity to flow speed (consistent with Craig et al.’s definition of *V* as speed), and *v*_0_ = 0.07 ≈ std(*X*) plays the role of *v*^*\**^. The ground-truth system is then:

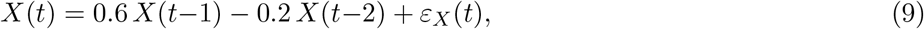

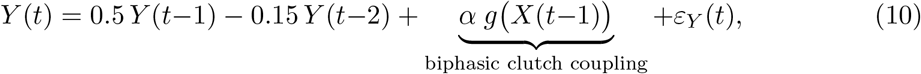

where *ε*_*X*_, *ε*_*Y*_ ∼ *N* (0, 0.05^2^). The clutch function *g*(*x*) is *even*: *g*(*x*) = *g*(−*x*), so positive and negative *X* produce identical coupling. This destroys any monotonic relationship between *X* and *Y* : Pearson, Spearman, and GC are expected to remain at chance level across all coupling strengths. CCM, operating on the reconstructed attractor, reads the deterministic relationship between *Y* ‘s manifold geometry and *X*’s state regardless of the sign symmetry.

#### (3c) Shared non-stationarity

In systems 1–3b, coupling is injected into the stochastic AR dynamics (*α X*(*t*−1) in system 1, or a nonlinear function of *X*(*t*−1) in systems 2 and 3b). In biological systems, however, coupling often manifests through shared deterministic components: cells migrating with correlated velocity trends, or proteins oscillating in phase. To test whether methods can detect coupling that resides entirely in deterministic structure, we construct two independent stochastic processes with no cross-coupling:

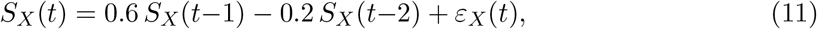

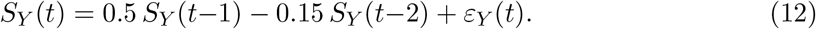

What we observe is each process plus shared deterministic components:

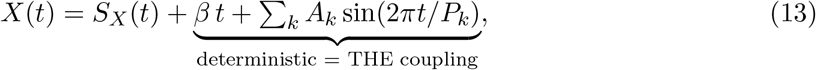

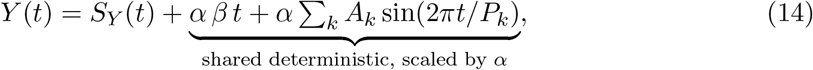

where *β* is the trend slope, *A*_*k*_ and *P*_*k*_ are cycle amplitudes and periods, and *α* controls how strongly *Y* ‘s deterministic components track those of *X*. At *α* = 0 the signals are fully independent; at *α* = 1 they share identical trends and cycles. This system is the mirror image of system 3a: in 3a, the biology is in the stochastic dynamics and the non-stationarity is an artifact; in 3c, the biology *is* the non-stationarity. Because the Stationarizer removes the trend and cycles, Granger causality operates on the independent residuals *S*_*X*_ and *S*_*Y*_ and is expected to detect nothing. CCM, operating on raw signals, preserves the correlated deterministic structure and is expected to detect the coupling. This system tests a coupling mode that is arguably the most common in biological time series yet is invisible to any method that requires stationarization.

#### (3d) Time-varying coupling strength

In biological systems, the strength of coupling between signals is rarely constant: adhesion maturation modulates the mechanical link between actin flow and traction force, cell cycle phase alters signaling responsiveness, and migration state determines whether neighboring cells interact. To model this, we use the same AR(2) backbone as system 1, but with coupling strength that oscillates sinusoidally over time:

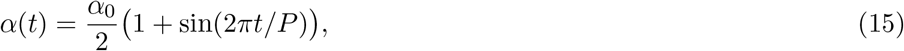

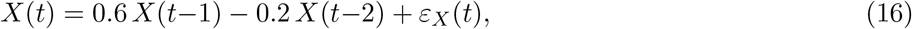

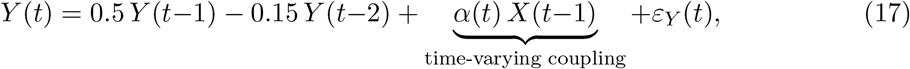

where *P* is the coupling oscillation period and *α*_0_ is the peak coupling strength (swept as *α*). The coupling cycles between zero and *α*_0_, so *X* drives *Y* only intermittently. Both *X* and *Y* are individually stationary—the non-stationarity resides exclusively in the coupling itself. Pearson and Spearman, which compute a single global correlation over the entire signal, average over coupled and uncoupled windows, diluting the statistic. CCM reconstructs the attractor from the coupled windows, where *Y* ‘s manifold encodes *X*’s state, and is expected to maintain high detection. The Stationarizer cannot remove this non-stationarity because the individual signals pass stationarity tests; only the relationship between them varies. Parameters: *P* = 100, *ε*_*X*_, *ε*_*Y*_ ∼ *N* (0, *σ*^2^=0.09), *n* = 300.

### Robustness analysis

#### (1) Sensitivity experiments

Beyond the ground truth systems above, we systematically varied signal length (*n* ∈ {20, 50, 100, 200}) and observation noise (SNR ∈ {∞, 20, 10, 5, 3, 0} dB, additive Gaussian), and compared four surrogate generation methods (IAAFT, shuffle, circular shift, block bootstrap with block size *b* = 10). These sweeps were run on three representative systems spanning the trichotomy— linear AR (system 1), biphasic clutch (system 3b), and shared non-stationarity (system 3c)—each at both null and coupled strengths. All experiments used 100 independent pairs per condition and 1000 IAAFT surrogates unless otherwise stated. Full experimental protocols are provided in S8. Ground truth systems: equations and experimental design.

#### (2) Shared stochastic driver (confounding control)

To test each method’s susceptibility to confounding, a latent variable *Z* independently drives both *X* and *Y* without direct interaction between them:

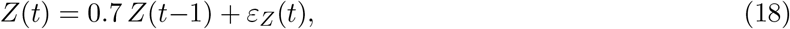

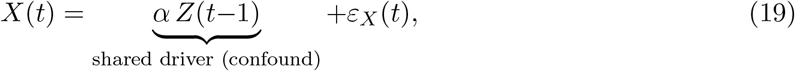

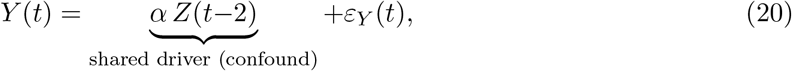

where *ε*_*Z*_, *ε*_*X*_, *ε*_*Y*_ ∼ *N* (0, *σ*^2^=0.09). The ground truth is that *X* and *Y* are not directly coupled: any detected association is a false positive due to the shared driver *Z*. This system tests each method’s susceptibility to the confounding problem [8].

#### Computational details

All ground truth experiments were executed on a workstation equipped with a 6-core Intel Xeon w3-2425 (12 threads, 32 GB RAM) with CPU-level pair parallelism via joblib (10 workers). The full benchmarking suite (ground truth systems across nine coupling strengths × 100 pairs, plus sensitivity sweeps across length, noise, and surrogate type) totaled tens of hours of computation. The NK cell application (878,879 pairs across 13 movies, up to 700 IAAFT surrogates per pair with sequential early stopping) was executed on the same workstation with 10 parallel workers in 29.8 h, using a vectorized CPU implementation of CCM that reproduces the reference results exactly (validated on 100 pairs). GPU timings on an NVIDIA A100 are reported in Table 5.

### Coupling detection methods

Signal Associa implements six association methods spanning three methodological families: linear correlation (Pearson, Spearman), information-theoretic dependence (mutual information, transfer entropy), and model-based causality (Granger causality, convergent cross-mapping). Each method targets a different subset of the coupling regimes defined above, and no single method reliably detects all four (Table 4). Below we describe each method and its key equation; detailed mathematical formulations are provided in S1–S5 Texts.

**Table 4.** Default configuration for each association method in Signal Associa. Signal version indicates whether the method receives raw or stationarized signals. *N*_surr_ indicates the default number of IAAFT surrogates for empirical p-value computation. Methods with *N*_surr_ = 0 use parametric p-values.

| Method | Measures | Signal version | $N_{\text{surr}}$ |
| --- | --- | --- | --- |
| Pearson correlation | Linear co-variation | Raw | 0 |
| Spearman correlation | Monotonic co-variation | Raw | 0 |
| Mutual information | Arbitrary dependence | Raw | 100 |
| Granger causality | Linear predictive causality | Stationary | 0 |
| CCM | Nonlinear deterministic causality | Raw | 100 |
| Transfer entropy | Directed information flow | Raw | 100 |

#### (1) Pearson and Spearman correlation

Pearson’s correlation coefficient *r* measures the strength of the linear relationship between two signals:

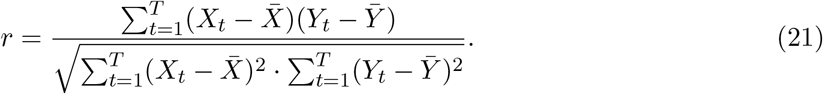

Spearman’s *ρ*_*s*_ replaces values with ranks before computing *r*, capturing monotonic nonlinear relationships. Both are symmetric and instantaneous: they cannot determine causal direction or detect lagged coupling. Signal Associa computes parametric p-values via the *t*-distribution (see S5. Correlation methods: Pearson and Spearman).

#### (1b) Lag-scanned Pearson (Danuser baseline)

As a baseline representing standard practice in cell biology [11], we also implement lag-scanned Pearson cross-correlation. The correlation is computed across a range of time lags *k* ∈ {−15, …, +15},

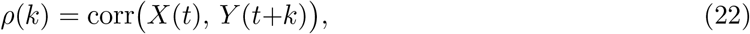

and the signed extremum *ρ*^*\**^ = *ρ*(*k*^*\**^) with *k*^*\**^ = arg max_*k*_|*ρ*(*k*)| is taken as the test statistic (the significant maximum/minimum of Machacek & Danuser [11]). Significance is assessed two ways: (i) the parametric 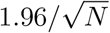 confidence level for Pearson’s coefficient used in the original method, and (ii) IAAFT surrogate testing. This baseline isolates the effect of lag scanning and of the parametric threshold, and is compared against the surrogate-controlled methods throughout the ground truth benchmarks.

#### (2) Mutual information

Mutual information quantifies any statistical dependence between two signals, including nonlinear and non-monotonic relationships [35]:

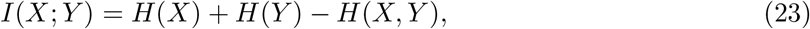

where *H*(·) denotes Shannon entropy. *I*(*X*; *Y* ) = 0 if and only if *X* and *Y* are statistically independent. Signal Associa estimates MI using the *k*-nearest neighbor method of Kraskov et al. [36] and reports normalized MI (NMI ∈ [0, 1]). Like correlation, MI is symmetric and cannot determine causal direction. Because MI has no simple parametric null distribution, surrogate testing is always used for significance assessment (see S4. Mutual information: nonlinear dependency detection).

#### (3) Granger causality

Granger causality tests whether the past of signal *X* improves prediction of *Y* beyond what *Y* ‘s own past provides [16, 35]. The null hypothesis is tested by comparing a restricted model

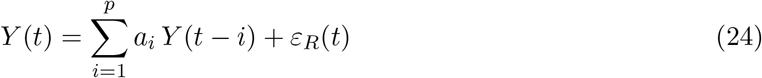

against an unrestricted model

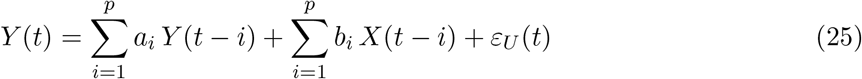

via the *F* -statistic. Signal Associa selects the lag order *p* by minimizing the Bayesian Information Criterion [37] (up to *p* = 5). Granger causality is directional and requires stationarity: Signal Associa automatically applies stationarization before this method while leaving other methods to operate on raw signals. A wild bootstrap [38] provides robust p-values for short signals (see S1. Granger causality: VAR models and bootstrap inference).

#### (4) Convergent cross-mapping

CCM tests whether *Y* ‘s state-space manifold contains recoverable information about *X*, which occurs when *X* causally influences *Y* [3]. The method exploits Takens’ embedding theorem [39]: *Y* is embedded into a delay-coordinate vector

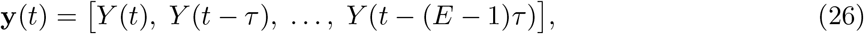

and nearest neighbors in this reconstructed manifold are used to cross-predict the corresponding values of *X*. The cross-map correlation

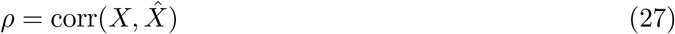

quantifies coupling strength. Signal Associa implements CCM with leave-one-out cross-validation, exponential distance weighting, and lag scanning across *l* ∈ {−2, −1, 0, 1, 2}, matching the reference pyEDM implementation to floating-point precision (max |Δ*ρ*| = 3.5 × 10^*−*7^ ; S7. CCM validation against pyEDM). CCM is directional, operates on raw signals, and is the only method in Signal Associa designed for nonlinear deterministic coupling (see S2. Convergent cross-mapping: state-space reconstruction).

#### (5) Transfer entropy

Transfer entropy measures the directed information flow from *X* to *Y* : the reduction in uncertainty about *Y* ‘s future when *X*’s past is known, beyond what *Y* ‘s own past provides [40]:

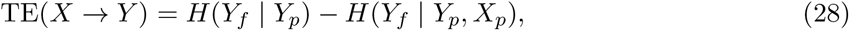

where *Y*_*f*_ is the future of *Y, Y*_*p*_ its past, and *X*_*p*_ the past of *X*. For Gaussian processes, TE is mathematically equivalent to Granger causality [41]; for nonlinear or non-Gaussian processes, TE captures dependencies that linear methods miss. Signal Associa estimates TE using *k*-nearest neighbor density estimation and reports both directional TE values and the net transfer entropy TE_net_ = TE(*X* → *Y* ) − TE(*Y* → *X*) (see S3. Transfer entropy: information-theoretic coupling).

#### (6) Stationarity and the Stationarizer

Among the six methods above, Granger causality (Eqs. 24–25) requires stationary inputs; applying it to signals with trends or cycles produces spurious detections. The remaining five methods operate on raw signals. Signal Associa addresses this asymmetry through an automatic Stationarizer that decomposes each signal as

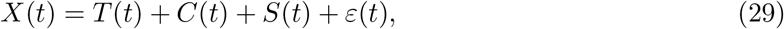

and removes the trend *T* (*t*) and cyclic component *C*(*t*) while preserving the stochastic component *S*(*t*) that carries causally relevant fluctuations.

A signal is weakly (covariance) stationary when three conditions hold simultaneously:

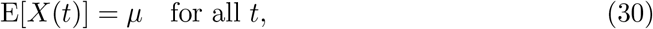

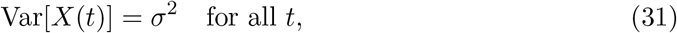

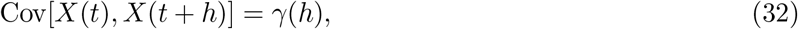

where Eq. 30 requires constant mean, Eq. 31 requires constant variance, and Eq. 32 requires that autocovariance depends only on the lag *h*, not on absolute time. The Stationarizer verifies these through a three-test diagnostic battery: the Augmented Dickey-Fuller (ADF) test [42] (null: unit root), the KPSS test [43] (null: stationary, complementary to ADF), and the Phillips-Perron test [44] (null: unit root, robust to heteroscedasticity). Condition (ii) is additionally assessed via a rolling variance ratio. A signal receives “high” stationarity confidence when all tests pass and variance is stable, “moderate” when at least two pass, and “low” otherwise.

Six stationarization methods are available, ordered by how much biological signal they preserve:

##### Bandpass filtering

Zero-phase Butterworth filter (default passband 0.008–0.25 Hz) [45]. Removes both low-frequency trends and high-frequency noise while preserving the coupling-relevant mid-frequency band. This is the gentlest method and the default first step.

##### Polynomial detrending

Fits a linear or quadratic polynomial to the signal and subtracts it. The degree is selected automatically: quadratic is used only when it improves *R*^2^ over linear by more than 0.05. Effective for monotonic drifts such as photobleaching.

##### Lazy cycle separation

A Takens embedding-based decomposition [46, 47] that isolates deterministic cycles from the stochastic component. Designed for signals with strong periodic content (e.g., oscillatory gene expression) where bandpass filtering would also remove coupling information near the cycle frequency.

##### Differencing

First-order differencing [48], Δ*X*(*t*) = *X*(*t*) − *X*(*t* − 1), removes unit roots and linear trends but amplifies high-frequency noise. Used as a last resort when gentler methods fail to achieve stationarity.

##### Combination methods

Two two-stage methods are available for signals where a single transformation is insufficient. In *bandpass + difference*, the signal is first bandpass-filtered, then the diagnostic battery is re-evaluated; differencing is applied to the filtered output only if it remains non-stationary. *Lazy + difference* follows the same logic: lazy cycle separation is applied first, and differencing is added only if the extracted stochastic component still fails the stationarity tests. In both cases, the gentler method handles the dominant non-stationarity (trends or cycles), and differencing addresses any residual unit root.

##### Automatic mode

The default mode (auto) applies these steps sequentially following the decomposition in Eq. 29: first check stationarity, then detrend if a polynomial trend explains significant variance (*R*^2^ *>* 0.05), then remove up to three dominant spectral peaks via FFT if any individual peak exceeds 5% of total spectral power, and finally difference as a last resort. Stationarity is re-evaluated after each step, and the pipeline stops as soon as the diagnostic battery passes. A spectral concentration metric (fraction of total power in the top two FFT bins, threshold 0.50) supplements the ADF/KPSS/PP battery to detect deterministic cycles that unit-root tests miss (Fig S6).

### Validation framework

#### Surrogate-based significance testing

Coupling statistics such as mutual information, convergent cross-mapping, and transfer entropy produce continuous-valued scores for which no standard parametric null distribution exists. A high score alone does not establish significance: biological time series possess intrinsic autocorrelation and spectral structure that can produce spurious coupling scores even between independent signals. Signal Associa addresses this through surrogate-based significance testing [19], where the observed coupling is compared against an empirical null distribution constructed from phase-randomized or permuted versions of the signal.

The procedure operates as follows. First, the association statistic *s*_real_ is computed on the original signal pair (*X, Y* ). Second, *N* surrogate versions of the predictor signal *X* are generated, each preserving certain statistical properties of *X* while destroying the coupling structure between *X* and *Y* . Third, the same association statistic is computed for each surrogate pair 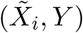, yielding a null distribution {*s*_1_, *s*_2_, …, *s*_*N*_ }. The empirical p-value quantifying the significance of the observed coupling is then computed as [49]:

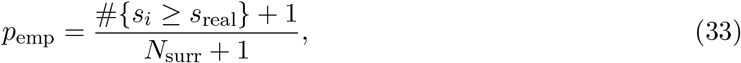

where *N*_surr_ is the number of surrogates. The +1 corrections in both numerator and denominator prevent *p* = 0 and ensure a valid p-value even when no surrogate exceeds the real statistic. A significance threshold of *p <* 0.05 is applied by default. Effect size is computed as

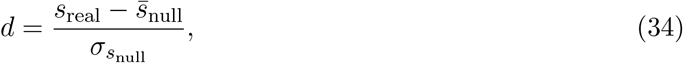

analogous to Cohen’s *d*, quantifying how many standard deviations the real statistic lies from the null distribution mean. An effect size *d >* 2 indicates that the observed coupling is far outside the range expected under the null hypothesis, providing confidence in the result independent of the p-value.

#### Surrogate methods

The choice of surrogate method determines the null hypothesis being tested, because different methods preserve different signal properties. Signal Associa implements four surrogate methods, ordered by the strength of the null hypothesis they impose:

1. **IAAFT (Iterated Amplitude-Adjusted Fourier Transform)**. The gold standard for autocorrelated signals [50]. Preserves both the amplitude distribution and the power spectrum (autocorrelation structure) of the original signal while randomizing the Fourier phases, thereby destroying cross-correlation with any other signal. The algorithm iterates between rank-matching to the original amplitude distribution and replacing Fourier magnitudes with those of the original signal until convergence (typically 100 iterations). Because IAAFT surrogates retain the temporal autocorrelation of the original signal, they provide the most conservative null hypothesis: a coupling detected against IAAFT surrogates cannot be explained by the spectral properties of the individual signals alone.
2. **Shuffle**. Random permutation of the signal values. Destroys all temporal structure, providing the weakest null hypothesis. Appropriate only when the signal has no meaningful autocorrelation; for biological time series with inherent temporal structure, shuffle surrogates inflate the apparent significance of detected coupling.
3. **Circular shift**. Circular time shift by a random offset:

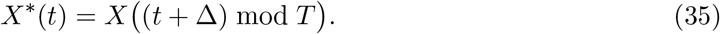 Preserves the full autocorrelation structure while destroying cross-correlation at specific lags. A good compromise for paired tests where preserving the temporal dynamics of each signal is important.
4. **Block bootstrap**. Resamples contiguous blocks of configurable length (default: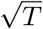 where *T* is signal length) [51]. Preserves short-range autocorrelation while disrupting long-range dependencies.

Signal Associa defaults to IAAFT surrogates for mutual information, CCM, and transfer entropy (Table 4). Pearson and Spearman correlations use parametric p-values, and Granger causality uses the F-test p-value from the VAR model, so neither requires surrogate testing by default.

#### GPU-accelerated computation

Modern biological experiments produce hundreds of simultaneously recorded trajectories, yielding tens to hundreds of thousands of signal pairs that each require surrogate-based significance testing. In the NK cell application, the 13 movies yield 878,879 signal pairs; with up to 700 surrogates per pair, this amounts to several hundred million CCM evaluations. Signal Associa addresses this through two complementary parallelization strategies: CPU-level pair parallelism (distributing independent signal pairs across cores) and GPU-level surrogate parallelism (generating and evaluating all *N* surrogates simultaneously via CuPy). We benchmarked both on the three initial NK movies (one per tumor line; 28,633 pairs, 700 surrogates per pair). On an NVIDIA A100 GPU, the analysis completed in 13.8 hours (1.7 seconds per pair on average), a 40.7× speedup over serial CPU computation, with stable per-pair timing across conditions (Table 5). For CPU-only systems, two further optimizations make large runs practical: a vectorized CCM implementation that shares the nearest-neighbor search across surrogates, and sequential early stopping, which halts surrogate testing once a pair’s significance decision is determined. With these, the full 13-movie analysis ran on a single 10-core workstation in 29.8 hours. Both implementations preserve algorithmic fidelity to the reference pyEDM package (max |Δ*ρ*| = 3.5 × 10^*−*7^; S7. CCM validation against pyEDM). Full implementation details are provided in S9. GPU-accelerated surrogate testing.

**Table 5.** Computational cost of CCM surrogate testing for the initial NK cell movies. (one movie per tumor line; 700 IAAFT surrogates per pair; A100 measured on Google Colab, 40 GB VRAM).

| Condition | Pairs | Per pair | Wall-clock |
| --- | --- | --- | --- |
| HT29 (MSS) | 15,324 | 1.71 s | 7.3 h |
| LoVo (MSI-H) | 6,699 | 1.79 s | 3.3 h |
| SW48 (MSI-H) | 6,610 | 1.67 s | 3.1 h |
| All conditions | 28,633 | 1.72 s (avg) | 13.8 h |
| CPU, serial (1 core) | 28,633 | $\sim 70$ s | $\sim 23$ days |
| CPU, parallel (10 cores) | 28,633 | $\sim 8$ s | $\sim 2.6$ days |
| *GPU speedup: $40.7\times$* |  |  |  |

### Biological data

#### Microfluidic chip and cell culture

NK cell migration data were acquired from a microphysiological tumor-vascular interface system [23]. The device is an injection-molded polystyrene chip in 384-well plate format containing three microchannels: a central hydrogel channel (100 *µ*m tall) flanked by two side channels (400 *µ*m tall), each approximately 1 mm wide and 4 mm long, sealed with a pressure-sensitive adhesive film and plasma-treated to ensure stable hydrogel patterning. Hydrogel was first injected into the central channel to create a patterned matrix. Cancer cells (∼0.5×10^6^ cells/mL) mixed with fibrin gel and co-seeded with HUVECs and fibroblasts were introduced into the central channel (Day 0); microtumors of approximately 100–200 *µ*m diameter formed after ∼5 days of culture. Endothelial cells were then seeded into the two side channels and cultured for approximately 5 days; the chip was tilted so that the endothelial cells attached to both walls where they contacted the hydrogel, forming a perfusable vessel. NK cells were loaded on Day 2 for live-cell imaging as they migrated through the vascular interface toward the tumor region.

#### Live-cell imaging

Four-channel fluorescence imaging was performed at 10× magnification (pixel size 2.4748 *µ*m) at 5-minute intervals for 273 frames (∼23 hours): NK cells (647 nm), cancer cells (488 nm), endothelial monolayer (405 nm), and dead cells (594 nm, Sytox). Three molecularly distinct colorectal cancer lines were used: HT29 (MSS, BRAF-mutant, low cytotoxicity), LoVo (MSI-H, KRAS-mutant, high cytotoxicity), and SW48 (MSI-H, KRAS/BRAF wild-type, high cytotoxicity). Thirteen movies were analyzed (HT29: 5, LoVo: 4, SW48: 4), each a separate field of view, acquired from 13 chips, one movie per chip, each chip from an independent experiment. After quality filtering (≥20 frames), 5,387 trajectories were retained for coupling analysis (HT29: 1,829, LoVo: 2,353, SW48: 1,205).

#### NK cell and tumor segmentation and tracking

Movies acquired as z-stacks (3 slices) were maximum-projected before analysis. NK cells (647 nm channel) and tumors (488 nm channel) were segmented with CellADE, our adaptive segmentation module. Each image was contrast-enhanced (CLAHE, clip limit 0.03); object centers were detected as Laplacian-of-Gaussian blobs (scale range *σ* = 2–8 pixels for NK cells and 10–20 pixels for tumors); foreground was separated around each center by local two-class fuzzy c-means clustering (membership threshold 0.4 for NK cells and 0.2 for tumors); and touching objects were split by seeded watershed. Mesh-like artifacts were removed using the ratio of an object’s total perimeter, including internal holes, to that of a circle of equal area (compact objects give about 1–1.5; objects with a ratio above 3 were discarded). NK cells were segmented in every frame; tumors, which were static over the recording, were segmented once per movie. NK cell centroids were linked into trajectories by linear assignment with gap closing (LapTrack): a maximum frame-to-frame displacement of 25 pixels and gap closing over up to 10 frames within 40 pixels. Trajectories shorter than 20 frames were excluded from coupling analysis. The tumor masks define the Keller–Segel field and the tumor regions used in the tumor-level analysis.

#### Signal extraction

Each NK cell trajectory is a bivariate time series (*x*(*t*), *y*(*t*)). To produce a univariate signal for coupling analysis, we project each trajectory onto its first principal component (PC_1_): the axis of maximum positional variance. This is the optimal linear compression of 2D motion into 1D, preserving the dominant kinematic mode while discarding the minor axis (typically lateral jitter). Alternative representations, Euclidean distance from origin (discards directional information), polar coordinates (yields two signals), speed (insensitive to coordinated turns), are each insufficient for coupling analysis. Across the 265 trajectories of the representative HT29 movie, PC_1_ captured 82 ± 11% of the total positional variance (range: 55–99%), confirming that NK cell motion is predominantly one-dimensional at the single-cell level.

#### Pairwise analysis

From each movie, all valid pairs (≥20 frames temporal overlap) were tested with CCM using up to 700 IAAFT surrogates (878,879 pairs across the 13 movies). Surrogates were evaluated in blocks of 10 with sequential early stopping [52]: a pair was stopped as soon as more surrogates exceeded its cross-map correlation than allowed for *p <* 0.05 at 700 surrogates, so significance decisions are identical to running all 700.

#### Network construction and metrics

For each movie, we constructed a coupling network from the CCM results. Each node represents one NK cell trajectory, and an edge connects cells *i* and *j* when their coupling is significant (*p <* 0.05 against the IAAFT surrogates); only pairs with ≥20 frames of temporal overlap are tested. Networks were restricted to migrating cells, defined by a track extent (the largest distance between any two positions of the track) of at least 100 *µ*m (84% of trajectories); this excludes cells that only fluctuate around one position. The resulting undirected graph *G* = (*V, E*) has *N* = |*V* | nodes. The degree *k*_*i*_ of a cell is the number of distinct cells it is coupled with.

We quantified network structure with four measures. *Overall coordination:* the significance rate *S* = |*E*| */* |*E*_tested_|, the fraction of tested pairs that show significant coupling in a movie. *Local coordination:* the clustering coefficient of node *i*, the fraction of its neighbors that are also connected to each other,

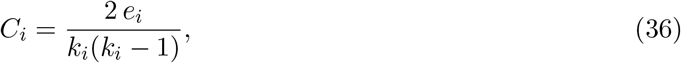

where *e*_*i*_ is the number of edges among the neighbors of *i*. To test whether clustering exceeded what network density alone produces, each movie’s network was compared with 5 degree-preserving randomizations (double-edge swaps, 2|*E*| swaps) and 5 random graphs with the same number of nodes and edges. *Global reach:* eigenvector centrality, which scores a cell by the centrality of the cells it is coupled with. For adjacency matrix *A* = {*a*_*ij*_} (*a*_*ij*_ = 1 for an edge, 0 otherwise),

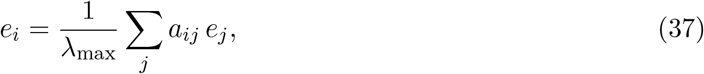

where *λ*_max_ is the largest eigenvalue of *A*, so **e** is the leading eigenvector of *A*. Because the definition is recursive, a cell scores high only if it is coupled to other high-scoring cells, so eigenvector centrality captures coordination beyond immediate neighbors. Because the unit-norm eigenvector scales as 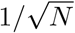, eigenvector centrality was multiplied by 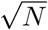 to make networks of different size comparable. *Relay role:* betweenness centrality, *b*_*i*_ = ∑_*s*≠*i*≠*t*_ *σ*_*st*_(*i*)*/σ*_*st*_, the fraction of shortest paths between other cells that pass through *i* (normalized; computed exactly). Because the network mean of normalized betweenness equals (*L* − 1)*/*(*N* − 2), with *L* the mean shortest-path length, betweenness was multiplied by (*N* − 2), as eigenvector centrality by 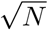, so that networks of different size are comparable. All metrics were computed on the unweighted graph, so they reflect network topology rather than CCM *ρ* magnitudes, using NetworkX (v3.1).

#### Tumor exposure and matched-exposure comparison

Tumor-derived signal was modeled as the steady state of the Keller–Segel chemoattractant equation, (*µ* − *D*_*C*_∇^2^) *C* = *S*, with *S* = 1 on tumor pixels, *D*_*C*_ = 100 px^2^, *µ* = 0.1 and *C* = 0 at the image border, solved on the pixel grid (decay length 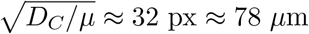). The exposure of an NK cell is the mean of |∇*C*| over the positions along its track. To compare tumor types at the same exposure (Fig 5a–c), the log_10_ exposures of all 4,538 network cells were divided into five quantile bins shared by all movies; the top bin, which holds the few cells sitting on tumor edges, was omitted. Within each bin, cells beyond 1.5 IQR of their tumor type were excluded, and each movie contributed one value, the mean metric of its cells in that bin (at least 5 cells). Lines show the mean ± s.e.m. over the movies of each type. The MSI-H minus HT29 difference was averaged over the four bins and tested by exact permutation: the 13 movies were reassigned to groups of 5 and 8 in all 1,287 possible ways, and *p* is the fraction of reassignments with an absolute difference at least as large as the observed one. The dependence of each metric on exposure within a type was summarized by Spearman’s *ρ* over that type’s cells. The same analysis with all cells in the networks, not only migrating ones, is reported in the text.

#### Tumor regions and NK presence

Each tumor contour was scaled by 1.3 about its centroid; pixels claimed by two tumors were assigned to the nearer one. For every tumor we counted the NK cell-frames inside this region over the movie, from all tracks including non-migrating cells, and expressed presence as cell-frames per unit area per frame. The network metric of a tumor is the mean over the network cells that entered its region, each weighted by the number of frames it spent there; tumors with fewer than 20 network cell-frames were excluded (262 tumors retained). Associations with presence (log_10_) were quantified by Spearman’s *ρ* and Theil–Sen fits over all tumors of a type (Fig 5d–f) and, as a check against differences in NK numbers between movies, within each movie, combined by Fisher-*z* averaging. Differences in *ρ* between tumor types were tested with Fisher’s *z*.

#### Recruitment over time

Each 22.7-h movie was divided into eleven windows of 2 h (24 frames). In each window and for each tumor we measured two quantities: the NK presence inside the tumor region, as above but over that window, and the integration of those NK cells, as the mean eigenvector centrality ×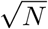 of the network cells that were inside the region during the window. For Fig 5g, each tumor was given one number, the centrality of its surrounding NK cells over the whole movie, and the tumors of each movie were ranked by it; the top third of each movie formed the high-centrality group and the bottom third the low-centrality group. For each group, the mean presence per window is plotted over time with the s.e.m. over tumors as the band.

#### Does integration predict recruitment?

We asked whether the integration of the NK cells around a tumor at one moment predicts how many NK cells arrive next. For every tumor and window *w* we took three numbers: the presence in *w*, the surrounding centrality in *w*, and the change in presence from *w* to a later window (*w* + 1, *w* + 2 or *w* + 3, i.e. 2, 4 or 6 h later). Each variable was centred within its movie, so that movies with many or few NK cells do not drive the result, and standardized. We then fitted a linear mixed model predicting the change in presence from the centrality and the presence in *w*, with a random intercept per tumor because each tumor contributes several windows (statsmodels MixedLM, REML). Including the presence in *w* means that the centrality effect is measured between tumors that held the same number of NK cells at that moment. Fig 5h shows the 2-h data as partial residuals: both the change in presence and the centrality after removing what the presence in *w* explains, averaged per sextile of centrality, with the fitted line. Fig 5i reports the standardized coefficient of centrality with its 95% confidence interval (Wald) for each tumor type and horizon; a positive value means that, at equal current presence, tumors with a more integrated NK network gained more NK cells afterwards.

#### Focal adhesion protein intensity data

Vinculin and FAK phosphorylation dynamics were co-imaged at individual focal adhesions using a spinning disk confocal microscope with a 60× objective (pixel size 108 nm). Three channels were acquired per timepoint: EGFP donor (ex 485/em 530 nm), FRET (ex 485/em 620 nm), and vinculin via SNAP-tag SNAP-Cell 647-SiR (ex 645/em 661 nm). FAK Y397 phosphorylation was reported by a Lyn-FAK FRET biosensor, where low FRET (high EGFP/FRET ratio) indicates active FAK [53]. Talin tension was reported indirectly through vinculin recruitment, as vinculin binds talin only when its rod domain unfolds under mechanical force. Images were acquired at 10-second intervals for 15 minutes (90 frames). Focal adhesions were detected and tracked from the vinculin channel using FASTIA, and the EGFP/FRET biosensor ratio was read along each adhesion trajectory, yielding paired vinculin(*t*) and pFAK(*t*) time series at individual adhesions. Cells were plated on polyacrylamide substrates of two stiffnesses: soft (0.6 kPa) and stiff (6 kPa).

Coupling between vinculin(*t*) and pFAK(*t*) was tested at each adhesion tracked for at least 15 frames, with CCM and lag-scanned Pearson against 700 IAAFT surrogates; adhesions whose trajectories were long enough for CCM (“tested” adhesions; 146 soft, 313 stiff) form the denominator of all rates. To define cell regions, a cell mask was obtained from the union of adhesion labels over time (morphological closing and hole filling), and a ∼12 *µ*m band inside the cell edge was divided by angle around the cell centroid into protruding, retracting and steady sectors, identified from the cell’s motion over the movie; adhesions in the cell interior were excluded from regional analyses. Coupling strength was compared with Mann-Whitney tests (one-sided for protruding versus other regions), detection rates with Fisher’s exact and *χ*^2^ tests, and directionality with binomial tests; per-region rates are shown with Wilson 95% confidence intervals.

### Software availability

Signal Associa is implemented in Python and is distributed as a module within CelLAB (Cellular Analysis Workbench), an open-source desktop platform for quantitative cell biology image analysis [54]. CelLAB follows the workbench paradigm of engineering software, providing a unified environment where specialized analysis pipelines share a common project system, interactive GUI, and reusable computational engines. Signal Associa requires NumPy, SciPy, and optionally CuPy for GPU-accelerated surrogate generation. Source code is freely available at https://github.com/HanLab-BME-MTU/CelLAB under the GNU General Public License v3.0. All code to reproduce the analyses in this paper, including ground truth benchmarking scripts and figure generation, is available at Zenodo (DOI: https://doi.org/10.5281/zenodo.23423612).

## Supporting information

Detailed mathematical formulations, algorithmic descriptions, validation results, and ground truth experimental designs are provided in Supporting Information (S1–S9).

## Acknowledgments

This work was supported by the Health Research Institute (HRI) at Michigan Technological University and by the National Institutes of Health (NIH) grant 2R15GM135806. We acknowledge the administrative support from the SOFT Foundry Institute at Seoul National University. This work was supported by the National Research Foundation of Korea (NRF) grant funded by the Korea government (MSIT) (RS-2021-NR059924, RS-2026-25522634), and the Ministry of Trade, Industry and Energy (MOTIE) and Korea Institute for Advancement of Technology (KIAT) through the International Collaborative R&D Program (P0030236).

## Supporting Information

### S1. Granger causality: VAR models and bootstrap inference

Granger causality fundamentally tests whether incorporating the historical data of one time series (*X*) improves the prediction of another time series (*Y* ) beyond the predictive power achievable using only the past values of *Y* itself. The evaluation involves comparing a restricted model (predicting *Y* from its own past only) against an unrestricted model (predicting *Y* from the past of both *Y* and *X*).

Consider two time series (*Y*_*t*_, *X*_*t*_) observed at *t* = 1, …, *T* . The Vector Autoregressive (VAR) model describes their interrelationships:

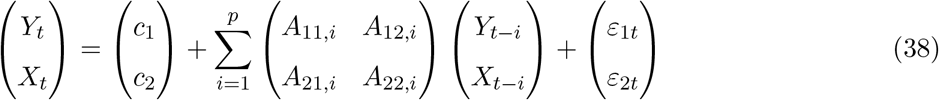

where *A*_12,*i*_ quantifies the impact of the *i*-th lagged value of *X* on the current value of *Y* and is central to the causality test.

The null hypothesis *H*_0_ : *A*_12,1_ = *A*_12,2_ = … = *A*_12,*p*_ = 0 (no Granger causality from *X* to *Y* ) is tested via the F-statistic:

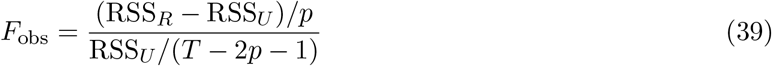

where RSS_*R*_ and RSS_*U*_ are the residual sums of squares from the restricted and unrestricted models, respectively.

The lag order *p* is selected by minimizing the Bayesian Information Criterion [37]:

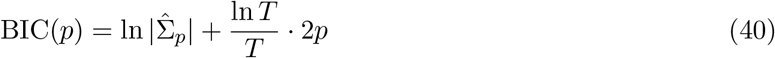

For robust inference, Signal Associa implements a wild bootstrap procedure [38]. Bootstrap samples are generated under *H*_0_ by setting *A*_12,*i*_ = 0 and resampling residuals:

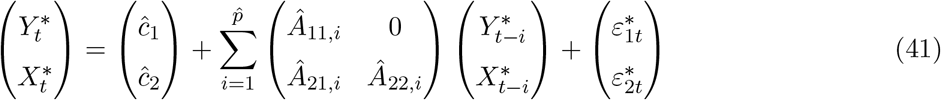

The bootstrap p-value is computed as *p*_boot_ = *B*^*−*1^ ‘E_*B*_ I(*F* ^*\**^ ≥ *F*_obs_).

#### Stationarity requirement

Granger causality assumes that the underlying VAR process is *stationary*, the statistical properties (mean, variance, autocorrelation) do not change over time. When applied to non-stationary signals (e.g., with trends, drift, or changing variance), the *F* -test rejects the null hypothesis too frequently, producing spurious causality findings. This is because shared trends create apparent predictive relationships that disappear once the trend is removed.

Signal Associa addresses this by automatically stationarizing signals before Granger causality analysis while leaving other methods (CCM, correlation) to operate on raw data where appropriate. The stationarization pipeline is configurable (see S6 Text) and includes detrending, bandpass filtering, and differencing options.

#### Practical considerations for short signals

Biological time series are often short (*T <* 100 time points), which limits the maximum lag order *p* and reduces statistical power. Signal Associa uses BIC-based lag selection (Eq 40) with an upper bound of *p* ≤ 5 to avoid overfitting on short signals. The wild bootstrap provides more reliable p-values than the asymptotic *F* -distribution when *T* is small.

#### Numerical example

Consider two signals of length *T* = 100:

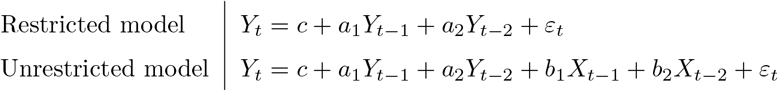

If *X* drives *Y*, the unrestricted model fits better: RSS_*U*_ *<* RSS_*R*_, yielding a large *F* . If the parametric *F* gives *p* = 0.003 and the bootstrap (700 resamples) gives *p*_boot_ = 0.004, both confirm that *X* Granger-causes *Y* at the 5% level.

#### Directionality

Granger causality is inherently directional: GC(*X* → *Y* ) and GC(*Y* → *X*) are computed independently by swapping the roles of *X* and *Y* in the VAR model. Signal Associa reports both directions and identifies the dominant one.

### S2. Convergent cross-mapping: state-space reconstruction

#### The question

We have two time series, *X* and *Y*, measured simultaneously (Fig S2a). We want to know: does *X* causally influence *Y* ? CCM answers this by testing whether *Y* ‘s dynamics contain information about *X*. The reasoning is that if *X* drives *Y*, then the history of *Y* should encode *X*’s influence, and we should be able to recover *X* from *Y* alone.

##### Step 1: Delay-embed *Y*

We take the *Y* time series and construct a set of vectors by pairing each value with its predecessor (Fig S2b):

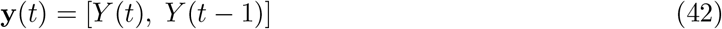

for embedding dimension *E* = 2 and time delay *τ* = 1. Each vector **y**(*t*) is a point in a two-dimensional space. The collection of all such points forms a cloud called the *reconstructed manifold* of *Y* .

Critically, this cloud contains no information about time ordering. Each point is simply a pair of numbers representing consecutive *Y* values. However, we retain a lookup table that records which time index *t* produced each point.

Consider 12 time points:

| $t$ | 1 | 2 | 3 | 4 | 5 | 6 | 7 | 8 | 9 | 10 | 11 | 12 |
| --- | --- | --- | --- | --- | --- | --- | --- | --- | --- | --- | --- | --- |
| $X(t)$ | 0.7 | 0.6 | 0.3 | 0.5 | 0.8 | 0.4 | 0.2 | 0.7 | 0.6 | 0.5 | 0.3 | 0.5 |
| $Y(t)$ | 0.2 | 0.5 | 0.8 | 0.4 | 0.1 | 0.6 | 0.9 | 0.3 | 0.1 | 0.5 | 0.8 | 0.4 |

After embedding *Y*, we obtain:

After embedding $Y$ , we obtain: 1221
| Label | Time | Embedded point $[Y(t), Y(t-1)]$ |
| --- | --- | --- |
| A | $t = 2$ | $[0.5, 0.2]$ |
| B | $t = 3$ | $[0.8, 0.5]$ |
| C | $t = 4$ | $[0.4, 0.8]$ |
| D | $t = 5$ | $[0.1, 0.4]$ |
| E | $t = 6$ | $[0.6, 0.1]$ |
| F | $t = 7$ | $[0.9, 0.6]$ |
| G | $t = 8$ | $[0.3, 0.9]$ |
| H | $t = 9$ | $[0.1, 0.3]$ |
| I | $t = 10$ | $[0.5, 0.1]$ |
| J | $t = 11$ | $[0.8, 0.5]$ |
| K | $t = 12$ | $[0.4, 0.8]$ |

Notice that points B and J are nearly identical: *Y* went through the same consecutive dynamics at *t* = 3 and *t* = 11, even though these moments are far apart in time. The embedding captures recurring dynamical patterns regardless of when they occurred. This is analogous to a pendulum returning to the same position-velocity state on different orbits.

##### Step 2: Find nearest neighbors

For each point in the embedded cloud, we find its *k* = *E* + 1 = 3 closest neighbors by Euclidean distance (Fig S2c). The value *k* = *E* + 1 comes from simplex projection theory: in an *E*-dimensional space, *E* + 1 points define a simplex (three points form a triangle in 2D).

For point D at *t* = 5, with coordinates [0.1, 0.4]:

| Neighbor | Time | Coordinates | Distance from D |
| --- | --- | --- | --- |
| H | $t = 9$ | $[0.1, 0.3]$ | 0.10 |
| A | $t = 2$ | $[0.5, 0.2]$ | 0.45 |
| E | $t = 6$ | $[0.6, 0.1]$ | 0.58 |

These three points are the moments when *Y* behaved most similarly to how it behaved at *t* = 5. They may be far apart in time, but they are close in state space, meaning *Y* went through similar consecutive changes.

##### Step 3: Cross-map to *X*

This is the key step that gives CCM its name (Fig S2d). We do not analyze *X*’s own structure. Instead, we use the neighbor times identified in *Y* ‘s manifold to look up what *X* was doing at those same moments.

D’s neighbors came from times *t* = 9, 2, 6. We read the corresponding *X* values:

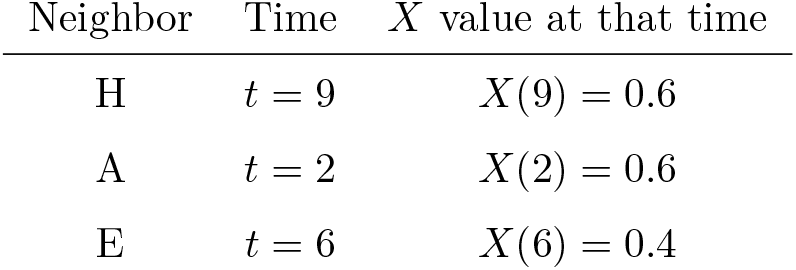

We compute exponential weights from the distances (closer neighbors contribute more):

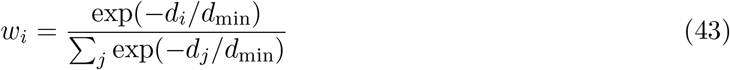

where *d*_min_ is the smallest neighbor distance. For our example:

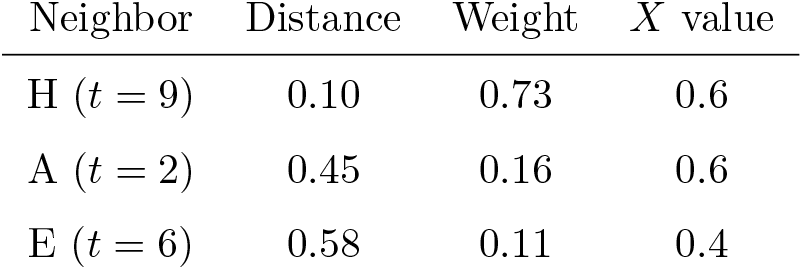

The predicted value of *X* at *t* = 5 is:

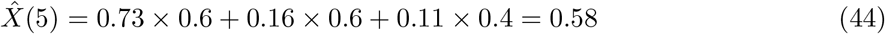

The actual value is *X*(5) = 0.8

##### Step 4: Repeat and evaluate

We repeat Steps 2–3 for every point in the embedded cloud (using leave-one-out: each point is predicted from its neighbors, excluding itself). This produces a vector of predicted values 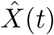 alongside the actual values *X*(*t*) (Fig S2e). The cross-map correlation is then computed across all points (Fig S2f):

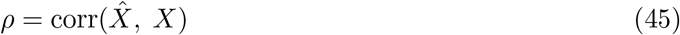

A high *ρ* means that *Y* ‘s manifold structure successfully recovers *X*’s values. This is only possible if *X*’s influence is encoded in *Y* ‘s dynamics, that is, if *X* causally drives *Y* .

A low *ρ* means *Y* ‘s dynamics contain no usable information about *X*, suggesting *X* does not drive *Y* .

##### The logic: why does this work?

The reasoning is rooted in Takens’ embedding theorem [39]. If *X* drives *Y*, then *Y* ‘s state at any moment reflects the history of *X*’s influence. When *Y* is in a particular dynamical state (a specific point in the embedded cloud), it arrived there because of a particular history of *X* values.

Therefore, when two points are close in *Y* ‘s manifold (meaning *Y* went through similar dynamics at both times), the *X* values at those times should also be similar, because similar *Y* states were caused by similar *X* inputs.

If *X* does not drive *Y*, there is no such correspondence, and *Y* ‘s neighbors will point to unrelated *X* values. The weighted average will be a poor prediction, and *ρ* will be low.

#### Directionality

CCM tests direction by running the procedure both ways. To test whether *X* drives *Y*, we embed *Y* and predict *X* (as above). To test whether *Y* drives *X*, we embed *X* and predict *Y* . If *X* drives *Y* but not vice versa, we expect *ρ*_*Y →X*_ (embed *Y*, predict *X*) to be higher than *ρ*_*X→Y*_ (embed *X*, predict *Y* ).

#### Significance testing

A single *ρ* value is difficult to interpret in isolation. Signal Associa tests significance by generating surrogate signals that preserve certain statistical properties of *X* (e.g., power spectrum via IAAFT) but destroy any causal relationship with *Y* . The CCM procedure is repeated for each surrogate, producing a null distribution of *ρ* values. The empirical p-value is the fraction of surrogate *ρ* values that equal or exceed the real *ρ*.

#### Parameters

- *E* (embedding dimension): the number of consecutive values used per vector. Default: *E* = 2.
- *τ* (time delay): the spacing between consecutive values in the embedding vector. Default: *τ* = 1.
- *k* (neighbors): set to *E* + 1.
- Lags: CCM is computed at multiple time lags *l* ∈ {−2, −1, 0, 1, 2}, and the best *ρ* is retained. This accounts for coupling delays.
- Library size: following the Sugihara et al. (2012) framework, prediction uses leave-one-out within the library rather than a held-out test set.

**Fig S2.**
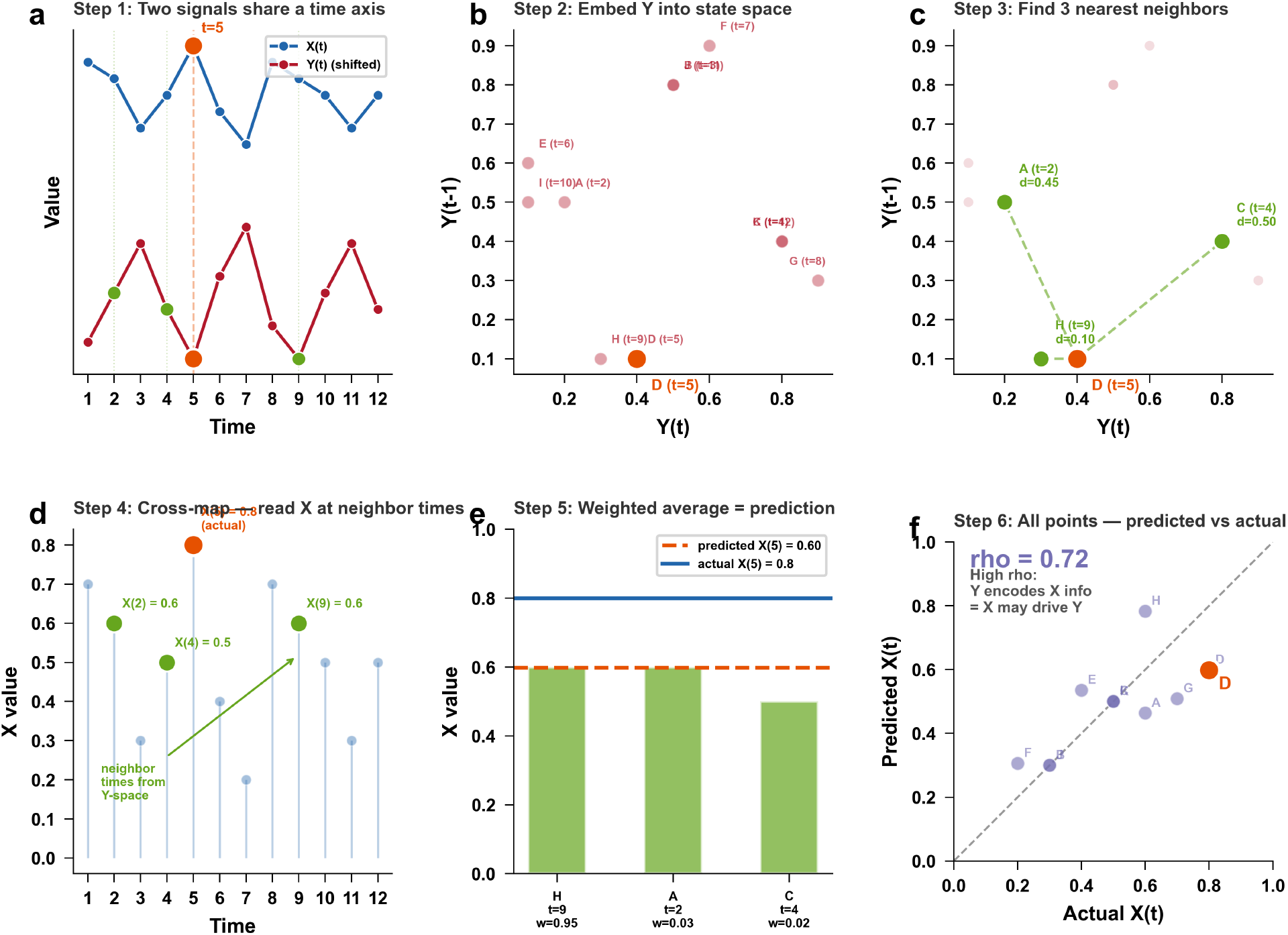
Step-by-step illustration of convergent cross mapping (CCM). (**a**) Two simultaneously recorded time series *X* and *Y* share a common time axis. Orange: target time point (*t* = 5); green: times identified as Y-neighbors in subsequent steps. (**b**) Delay embedding transforms *Y* into a cloud of state-space points [*Y* (*t*), *Y* (*t*−1)]. Each point represents consecutive *Y* dynamics; time ordering is lost, but time indices are retained as a lookup table. (**c**) For a target point (orange, *t* = 5), the *k* = 3 nearest neighbors are found by Euclidean distance in the embedded space. These are the moments when *Y* exhibited the most similar dynamics. (**d**) Cross-mapping: the neighbor times identified in *Y* ‘s manifold are used to look up the corresponding *X* values. This is the “cross” in cross-mapping: structure found in *Y*, values read from *X*. (**e**) A distance-weighted average of the neighbor *X* values produces the predicted 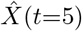 (orange dashed line). Comparison with the actual *X*(5) (blue solid line) evaluates prediction quality for this single point. (**f** ) Repeating Steps 2–5 for every embedded point yields a scatter of predicted versus actual *X* values. The Pearson correlation *ρ* quantifies how well *Y* ‘s manifold recovers *X*’s information: high *ρ* indicates *X* causally influences *Y* .

### S3. Transfer entropy: information-theoretic coupling

#### The question

Mutual information captures any dependence between *X* and *Y* but cannot determine direction. Transfer entropy (TE) adds directionality by asking: does knowing *X*’s past reduce uncertainty about *Y* ‘s future, *beyond what Y ‘s own past already provides*?

**Definition**. Transfer entropy from *X* to *Y* is defined as:

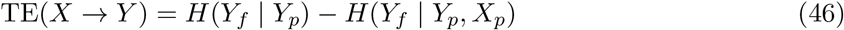

where *Y*_*f*_ is the future of *Y, Y*_*p*_ is the past of *Y*, and *X*_*p*_ is the past of *X*. The first term is the uncertainty about *Y* ‘s future given only *Y* ‘s own history. The second term is the uncertainty about *Y* ‘s future given both *Y* ‘s and *X*’s histories. The difference is the *additional* predictive information that *X* provides.

If *X* does not influence *Y*, knowing *X*’s past adds nothing: *H*(*Y*_*f*_ |*Y*_*p*_, *X*_*p*_) = *H*(*Y*_*f*_ |*Y*_*p*_) and TE = 0. If *X* drives *Y*, knowing *X*’s past reduces uncertainty about *Y* ‘s future and TE *>* 0.

#### Step-by-step numerical example

Consider 10 time points where *Y* tracks *X* with a one-step delay:

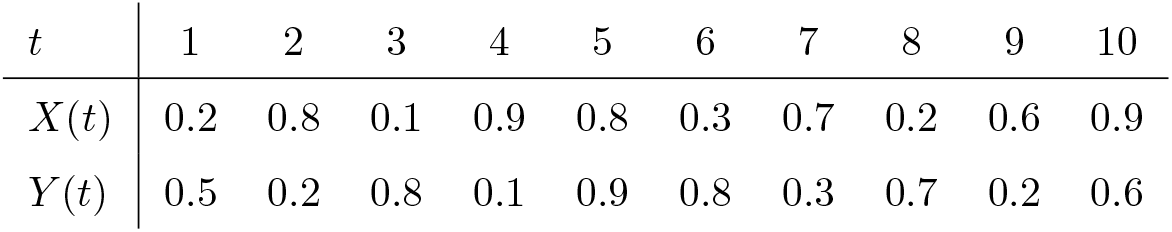

To compute TE by hand, we discretize each value as H (high, ≥ 0.5) or L (low, *<* 0.5) and use lag-1 histories.

##### Step 1: Build the triplets

For each time *t* = 2, …, 10, record (*Y*_*p*_, *X*_*p*_, *Y*_*f*_ ) where *Y*_*p*_ = *Y* (*t*−1), *X*_*p*_ = *X*(*t*−1), *Y*_*f*_ = *Y* (*t*):

| $t$ | $Y_p$ | $X_p$ | $Y_f$ |
| --- | --- | --- | --- |
| 2 | H | L | L |
| 3 | L | H | H |
| 4 | H | L | L |
| 5 | L | H | H |
| 6 | H | H | H |
| 7 | H | L | L |
| 8 | L | H | H |
| 9 | H | L | L |
| 10 | L | H | H |

##### Step 2: Compute *H*(*Y*_*f*_ |*Y*_*p*_), uncertainty using only *Y* ‘s past

When *Y*_*p*_ = H (5 occurrences): *Y*_*f*_ is L, L, H, L, L → *P* (L) = 4*/*5, *P* (H) = 1*/*5. 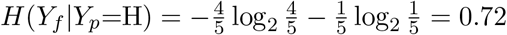 bits.

When *Y*_*p*_ = L (4 occurrences): *Y*_*f*_ is always H → *H*(*Y*_*f*_ |*Y*_*p*_=L) = 0.

Weighted average: 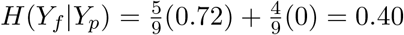 bits.

##### Step 3: Compute *H*(*Y*_*f*_ |*Y*_*p*_, *X*_*p*_), uncertainty using both histories

(*Y*_*p*_=H, *X*_*p*_=L): 4 cases, *Y*_*f*_ always L → *H* = 0. (*Y*_*p*_=L, *X*_*p*_=H): 4 cases, *Y*_*f*_ always H → *H* = 0. (*Y*_*p*_=H, *X*_*p*_=H): 1 case, *Y*_*f*_ = H → *H* = 0.

*H*(*Y*_*f*_ |*Y*_*p*_, *X*_*p*_) = 0, knowing both pasts eliminates all uncertainty.

##### Step 4: Transfer entropy

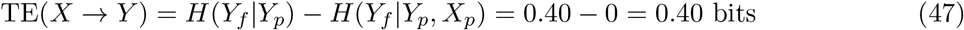

The past of *X* reduces uncertainty about *Y* ‘s future by 0.40 bits. When *Y*_*p*_ = H, *Y* ‘s own past leaves ambiguity (usually L, sometimes H), but adding *X*_*p*_ resolves it: if *X*_*p*_ = L then *Y*_*f*_ = L; if *X*_*p*_ = H then *Y*_*f*_ = H.

Running the same computation in the reverse direction yields TE(*Y* → *X*) = 0.10 bits, much smaller, confirming that the dominant information flow is from *X* to *Y* .

#### Relationship to Granger causality

For Gaussian processes, TE and Granger causality are mathematically equivalent (Barnett et al., 2009):

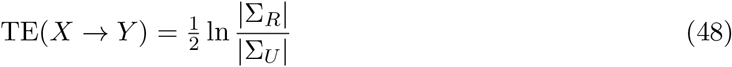

where Σ_*R*_ and Σ_*U*_ are the residual covariance matrices of the restricted and unrestricted VAR models. For non-Gaussian or nonlinear processes, TE captures dependencies that linear Granger causality misses.

#### Estimation

Signal Associa estimates TE using *k*-nearest neighbor density estimation in the joint space (*Y*_*f*_, *Y*_*p*_, *X*_*p*_) and marginal spaces, following the Kraskov–Stögbauer–Grassberger framework [36]. This avoids histogram binning and adapts to the local density of the data.

#### Net transfer entropy

Because TE can be non-zero in both directions even when only one causal link exists (due to shared dynamics or common driving), the *net* transfer entropy provides a cleaner directional measure:

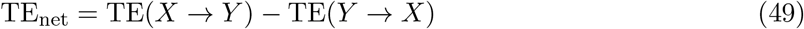

A positive value indicates dominant influence from *X* to *Y* .

#### Parameters

- Lag (*k*): number of past time steps considered. Default: *k* = 1.
- Prediction horizon: how far ahead *Y*_*f*_ is from *Y*_*p*_. Default: 1 step.
- *n*_neighbors_: number of neighbors for kNN density estimation. Default: 3.
- Surrogate testing is essential because TE has no simple parametric null distribution.

### S4. Mutual information: nonlinear dependency detection

#### The question

Pearson and Spearman correlation capture linear or monotonic relationships. But two signals can be strongly coupled in ways that are neither linear nor monotonic, for example, *Y* = sin(*X*) or *Y* = *X*^2^. Mutual information (MI) captures *any* statistical dependence, including nonlinear and non-monotonic relationships.

**Definition**. MI quantifies how much knowing one signal reduces uncertainty about the other:

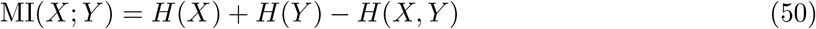

where *H*(*X*) is the Shannon entropy of *X, H*(*Y* ) of *Y*, and *H*(*X, Y* ) their joint entropy. Equivalently:

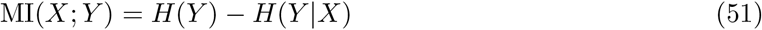

This reads: MI is the reduction in uncertainty about *Y* when *X* is known. If *X* and *Y* are independent, *H*(*Y* |*X*) = *H*(*Y* ) and MI = 0. If *Y* is perfectly determined by *X, H*(*Y* |*X*) = 0 and MI = *H*(*Y* ).

#### Numerical example

Consider 10 simultaneous measurements of two partially coupled

| $t$ | 1 | 2 | 3 | 4 | 5 | 6 | 7 | 8 | 9 | 10 |
| --- | --- | --- | --- | --- | --- | --- | --- | --- | --- | --- |
| $X(t)$ | 0.2 | 0.8 | 0.3 | 0.9 | 0.7 | 0.1 | 0.6 | 0.4 | 0.8 | 0.5 |
| $Y(t)$ | 0.3 | 0.7 | 0.4 | 0.8 | 0.6 | 0.2 | 0.9 | 0.6 | 0.3 | 0.7 |

To compute MI by hand, we discretize each value as H (high, ≥ 0.5) or L (low, *<* 0.5).

##### Step 1: Marginal distributions

To compute MI by hand, we discretize each value as H (high, $\geq 0.5$ ) or L (low, $< 0.5$ ). 1346 **Step 1: Marginal distributions.** 1348
| $X$ | Count | $P$ | $Y$ | Count | $P$ |
| --- | --- | --- | --- | --- | --- |
| L | 4 | 0.4 | L | 3 | 0.3 |
| H | 6 | 0.6 | H | 7 | 0.7 |

##### Step 2: Marginal entropies

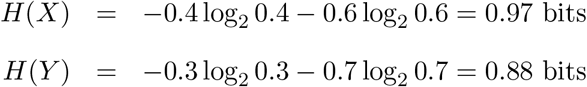

##### Step 3: Joint distribution

Counting how often each (*X, Y* ) combination occurs:

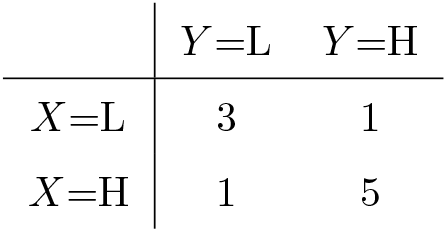

| | $Y=L$ | $Y=H$ |
| --- | --- | --- |
| $X=L$ | 3 | 1 |
| $X=H$ | 1 | 5 |

When *X* is low, *Y* is usually low (3 out of 4). When *X* is high, *Y* is usually high (5 out of 6). The signals are coupled but not perfectly, two mismatches exist (*t* = 8 and *t* = 9).

##### Step 4: Joint entropy

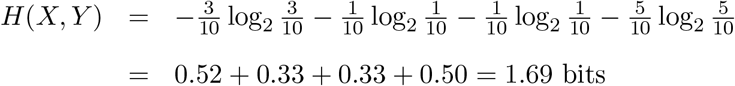

##### Step 5: Mutual information

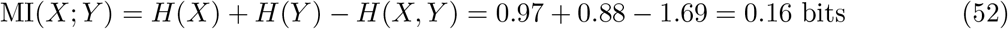

Interpretation: knowing *X* reduces uncertainty about *Y* by 0.16 bits. For comparison, if *X* and *Y* were independent, *H*(*X, Y* ) would equal *H*(*X*) + *H*(*Y* ) = 1.85 bits and MI would be zero.

##### Step 6: Normalized MI

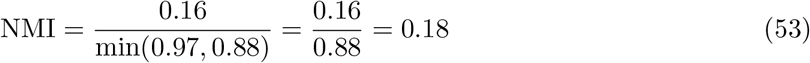

On a 0–1 scale, the coupling is weak to moderate, reflecting the two mismatched time points.

For continuous signals, MI is estimated using the *k*-nearest neighbor method of Kraskov, Stögbauer, and Grassberger [36], which avoids the information loss from discretization by estimating local densities from neighbor distances in the joint (*X, Y* ) and marginal spaces.

#### Symmetry and non-directionality

MI is symmetric: MI(*X*; *Y* ) = MI(*Y* ; *X*). It cannot distinguish cause from effect. This makes MI a measure of *association strength* rather than *causal direction*, complementary to the directional methods (GC, CCM, TE) in Signal Associa.

#### Normalization

Raw MI values depend on the entropy scales of the signals, making cross-comparison difficult. Signal Associa computes normalized MI:

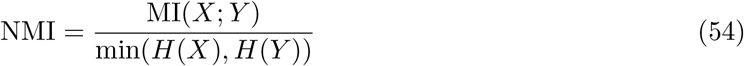

which ranges from 0 (independence) to 1 (complete dependence).

#### Why surrogate testing is essential

Unlike correlation, MI has no simple parametric null distribution. Two independent autocorrelated signals can produce non-zero MI purely from their temporal structure. Surrogate testing destroys the cross-signal relationship while preserving each signal’s autocorrelation, providing the correct null distribution for significance assessment.

### S5. Correlation methods: Pearson and Spearman

#### Pearson correlation

Pearson’s correlation coefficient *r* measures the strength of the *linear* relationship between two signals:

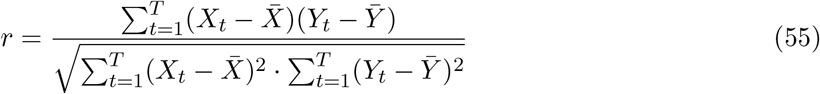

The value *r* ranges from −1 (perfect negative linear relationship) through 0 (no linear relationship) to +1 (perfect positive linear relationship). A parametric p-value can be obtained via the *t*-test 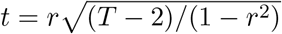 under the assumption that both signals are normally distributed and independent under *H*_0_.

#### Numerical example

Consider five time points:

| $t$ | 1 | 2 | 3 | 4 | 5 |
| --- | --- | --- | --- | --- | --- |
| $X(t)$ | 1.0 | 2.0 | 3.0 | 4.0 | 5.0 |
| $Y(t)$ | 2.1 | 3.9 | 6.2 | 7.8 | 10.1 |

Means: 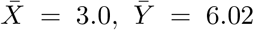. The deviations 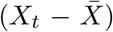 are (−2, −1, 0, 1, 2) and 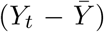 are (−3.92, −2.12, 0.18, 1.78, 4.08). The numerator is (−2)(−3.92) +(−1)(−2.12) + 0(0.18) +(1)(1.78) + (2)(4.08) = 20.0. After normalizing, *r* = 0.999, a near-perfect linear relationship.

#### Spearman correlation

Spearman’s *ρ*_*s*_ replaces each value with its rank and then computes Pearson’s *r* on the ranks:

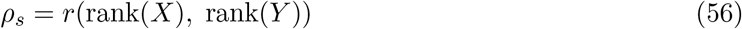

This captures *monotonic* relationships (consistently increasing or decreasing) even when the relationship is nonlinear. For example, if *Y* = *e*^*X*^, Pearson underestimates the association because the relationship curves, but Spearman returns *ρ*_*s*_ = 1.0 because the rank ordering is preserved perfectly.

#### Limitations

Both Pearson and Spearman are *symmetric*, they cannot distinguish whether *X* drives *Y* or *Y* drives *X*. They are also *instantaneous*, they compare *X*(*t*) to *Y* (*t*) at the same time point, missing lagged coupling entirely. A signal pair where *Y* (*t*) = *X*(*t* − 5) (perfect coupling with a 5-step delay) would yield *r* ≈ 0.

Furthermore, Pearson misses nonlinear coupling (e.g., *Y* = *X*^2^), and both methods are insensitive to dynamical coupling where the relationship is embedded in the state-space structure rather than point-by-point values.

#### When are surrogates needed?

The parametric *t*-test p-value assumes normality and independence, which biological time series routinely violate (autocorrelation inflates apparent significance). Signal Associa offers surrogate-based testing as an alternative: IAAFT surrogates preserve the autocorrelation structure of *X* while destroying any cross-signal coupling, providing a more conservative and realistic null distribution.

### S6. Stationarizer: preprocessing pipeline

The Stationarizer pipeline is described in Materials and methods. Fig S6 demonstrates the auto pipeline on four representative signal types and validates performance across 10 signal classes.

**Fig S6.**
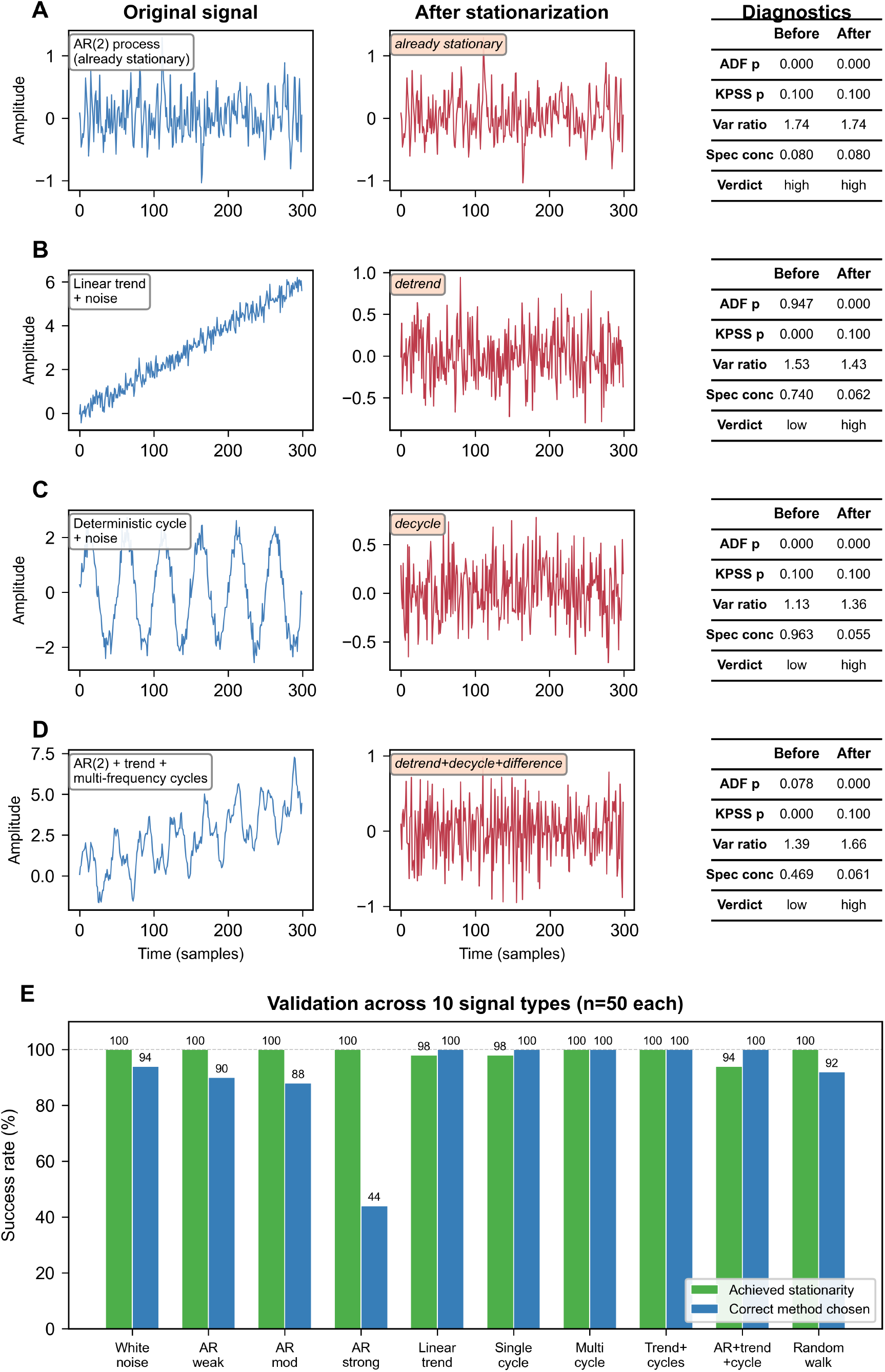
Stationarizer auto pipeline demonstration and validation. Panels A–D show four representative signal types (left: original signal, center: processed output, right: before-and-after diagnostics). (**A**) AR(2) process: correctly identified as already stationary. (**B**) Linear trend: removed by polynomial detrending. (**C**) Deterministic sinusoid: detected via spectral concentration and removed by FFT-based decycling. (**D**) Composite signal (AR + trend + two cycles): full pipeline (detrend + decycle + difference). (**E**) Validation across 10 signal types (*n* = 50 each). Green: achieved stationarity (≥94% all types). Blue: correct method chosen (≥88% except AR strong at 44%).

### S7. CCM validation against pyEDM

Signal Associa implements convergent cross-mapping from scratch to enable GPU-accelerated surrogate testing at scale. To confirm that the implementation produces results identical to the reference pyEDM package (Sugihara Lab, cppEDM v1.15.4), we conducted a systematic comparison across 10,000 independent CCM evaluations.

We generated 100 coupled autoregressive signal pairs (*n* = 300, coupling strength *α* = 0.4) and for each pair created 100 shuffle surrogates. Every surrogate was evaluated by three independent CCM engines: Signal Associa CPU (NumPy + SciPy cKDTree), Signal Associa GPU (CuPy), and pyEDM. All three received identical input arrays, used matching parameters (*E* = 2, *τ* = 1, leave-one-out cross-validation, exponential distance weighting), and computed bidirectional cross-map correlations.

Across all 10,000 comparisons, the maximum absolute difference between Signal Associa and pyEDM was 3.54 × 10^*−*7^ for both the CPU and GPU backends (Fig S7a,b). This discrepancy is consistent with differences in floating-point operation ordering between Python/SciPy and C++ implementations and confirms algorithmic equivalence. The CPU and GPU backends of Signal Associa agreed to machine precision (∼ 10^*−*14^; Fig S7c).

The validation establishes that Signal Associa’s CCM produces results indistinguishable from the reference implementation. Combined with surrogate-based significance testing (which pyEDM does not provide), the GPU-accelerated engine enables CCM analysis at scales infeasible with serial processing (Fig S7d).

**Fig S7.**
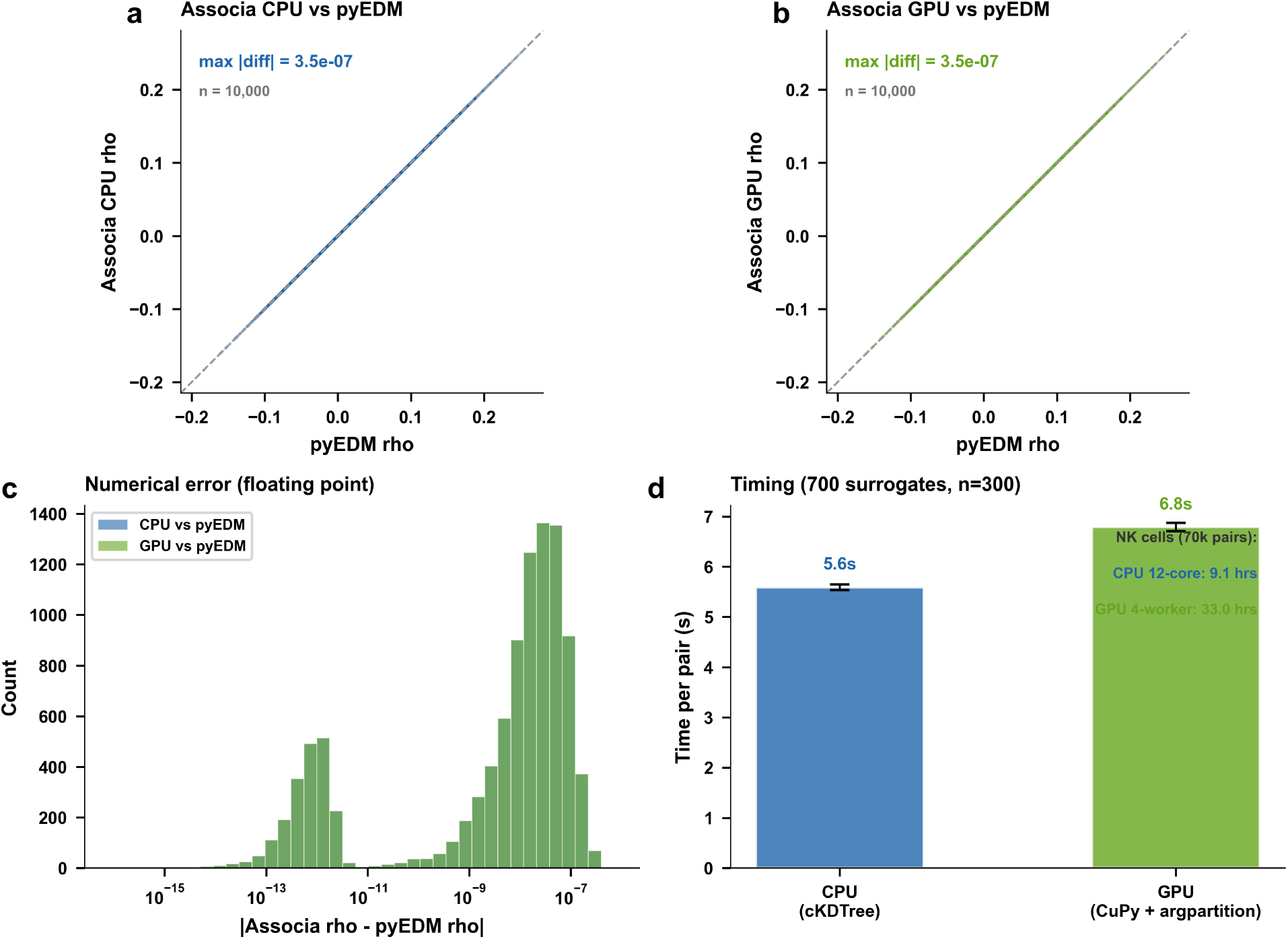
Validation of Signal Associa CCM implementation against pyEDM. (**a**) Scatter of cross-map correlation *ρ* computed by Signal Associa (CPU backend) versus pyEDM for 10,000 surrogate evaluations (100 coupled AR signal pairs × 100 shuffle surrogates each). Points fall on the identity line with a maximum absolute difference of 3.5 × 10^*−*7^, consistent with floating-point arithmetic ordering differences between Python/SciPy and C++ implementations. (**b**) Same comparison for the GPU backend (CuPy), showing identical precision to the CPU path. (**c**) Distribution of absolute numerical errors between Associa and pyEDM for both backends. All errors fall below 10^*−*6^, confirming algorithmic equivalence. (**d**) Computation time per signal pair (700 IAAFT surrogates, *n* = 300, *E* = 2, bidirectional with 5 lags) on a workstation equipped with a 12-core CPU and NVIDIA T1000 8 GB GPU. Text inset shows projected wall-clock time for the three initial NK cell movies (28,633 pairs).

### S8. Ground truth systems: equations and experimental design

This section documents the exact experimental protocols used in the ground truth bench-marking (Figs 2 and 3). System equations and coupling regimes are defined in Materials and methods. All parameter values, sweep ranges, and random seed conventions are specified to enable exact reproduction.

#### System-specific parameters

System 1 (coupled AR): *ε*_*X*_, *ε*_*Y*_ ∼ *N* (0, 0.09), 100 burn-in steps discarded, *n* = 300. System 2 (coupled logistic): *r*_*X*_ = 3.8, *r*_*Y*_ = 3.5 (chaotic regime), *β*_*XY*_ = 0.02 (fixed), initial conditions *X*(0) = 0.4 + *δ, Y* (0) = 0.4 + *δ* with *δ* ∼ *N* (0, 0.01), values clipped to [0.001, 0.999], 500 burn-in steps, *n* = 100.

#### Experiment 1: Coupling strength sweep (Fig 2)

The six ground truth systems and the shared-driver system are each swept across 9 coupling strengths. Systems 1, 2, 3a, 3c and 3d, and the shared-driver system 4 (driving strength), use

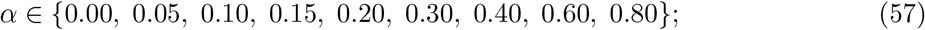

the clutch system 3b, whose bell-shaped coupling saturates at large inputs, uses

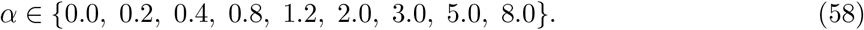

At each value, 100 independent signal pairs are generated (seeds 0–99) and tested with all six methods. The detection rate at *α* = 0 estimates the false positive rate (FPR).

#### Experiment 2: Directionality (Fig 2)

For each system and each *α >* 0, the ground truth is *X* → *Y* (unidirectional). Direction is classified as correct when the forward statistic exceeds the reverse statistic and the forward test is significant (*p <* 0.05). For symmetric methods (Pearson, Spearman, MI), directionality is undefined. For CCM, directionality uses unidirectional *ρ* (bidirectional=False).

#### Experiments 3–5: Sensitivity sweeps (Fig 3)

Three systems spanning the trichotomy are used: linear AR (system 1), biphasic clutch (system 3b) and shared non-stationarity (system 3c), each at a null (*α* = 0) and coupled strength, with 100 pairs per condition.

- *Signal length (Experiment 3): n* ∈ {20, 50, 100, 200}, with *α* ∈ {0, 0.15, 0.30, 0.60} (system 1), {0, 0.4, 1.2, 5.0} (system 3b) and {0, 0.05, 0.10, 0.20} (system 3c).
- *Observation noise (Experiment 4):* additive Gaussian noise at SNR ∈ {∞, 20, 10, 5, 3, 0} dB, where 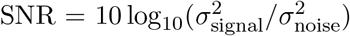, at *α* = 0.30 (system 1), 1.2 (system 3b) and 0.10 (system 3c), plus *α* = 0.
- *Surrogate method (Experiment 5):* IAAFT, random shuffle, circular shift and block bootstrap (block size *b* = 10), at the same coupled strengths as Experiment 4 plus *α* = 0, *n* = 300.

#### Common parameters across all experiments

- Surrogate count: *N* = 1000 (yields p-value resolution of 1*/*(1000 + 1) ≈ 0.001)
- Surrogate method: IAAFT with 100 iterations (except Experiment 5)
- Significance threshold: *p <* 0.05 (empirical p-value)
- CCM: *E* = 2, *τ* = 1, lag scan *l* ∈ {−2, −1, 0, 1, 2}, leave-one-out cross-validation (pyEDM-validated)
- Granger causality: BIC-selected lag order, *p*_max_ = 5
- Random seeds: pair *i* uses seed *i* for signal generation and surrogate testing (deterministic reproducibility)

### S9. GPU-accelerated surrogate testing

Computational cost and benchmark results are reported in Materials and methods (Table 5). This section describes the two parallelization strategies.

#### Strategy A: CPU pair-parallel

Independent signal pairs are distributed across CPU cores using joblib, with surrogates tested serially within each pair.

#### Strategy B: GPU surrogate-parallel

All *N* surrogates for a given pair are generated and tested simultaneously on the GPU via CuPy, proceeding in three stages that keep data on-device throughout: (1) *N* IAAFT surrogates are generated in parallel on an (*N* × *T* ) matrix with chunk-based memory management; (2) *k*NN distance computation, exponential weighting, and cross-map correlation are vectorized across all *N* surrogates in a single batched operation; (3) the null distribution is returned to CPU only after all *N* statistics are computed, eliminating repeated transfers.

Strategies A and B compose: pairs are distributed across CPU cores, each offloading surrogate testing to the GPU. Validation against pyEDM confirmed agreement to floating-point precision (max |Δ*ρ*| = 3.5 × 10^*−*7^; Fig S7).

### S10. Surrogate choice across all methods

The main-text comparison of surrogate methods (Fig 3g,h) focuses on CCM, the only method whose detection is strongly surrogate-dependent. Here we report detection and false-positive rates for all six methods under each surrogate type (IAAFT, shuffle, block bootstrap, and circular shift), across the AR, biphasic clutch, and shared non-stationarity systems (Fig S10).

Two patterns generalize across methods. First, on the shared non-stationarity null, shuffle surrogates inflated false positives for every autocorrelation-sensitive method, because shuffling destroys the temporal structure that a valid null must preserve: Pearson rose from 7% (IAAFT) to 18%, Spearman from 9% to 19%, CCM from 5% to 14%, and MI from 4% to 11% (Fig S10f). IAAFT and circular surrogates, which preserve autocorrelation, kept these rates near nominal. Second, block bootstrap selectively collapsed CCM detection (76% → 1% on AR, 96% → 8% on shared non-stationarity) while leaving the other methods unaffected, because preserving local temporal blocks retains the manifold structure that cross-mapping exploits and so removes the contrast against the null. Transfer entropy showed a mild baseline inflation (10–13% false positives) that was largely independent of surrogate type, reflecting its own estimator variance rather than the surrogate construction. These results confirm IAAFT as the best-calibrated default across methods.

**Fig S10.**
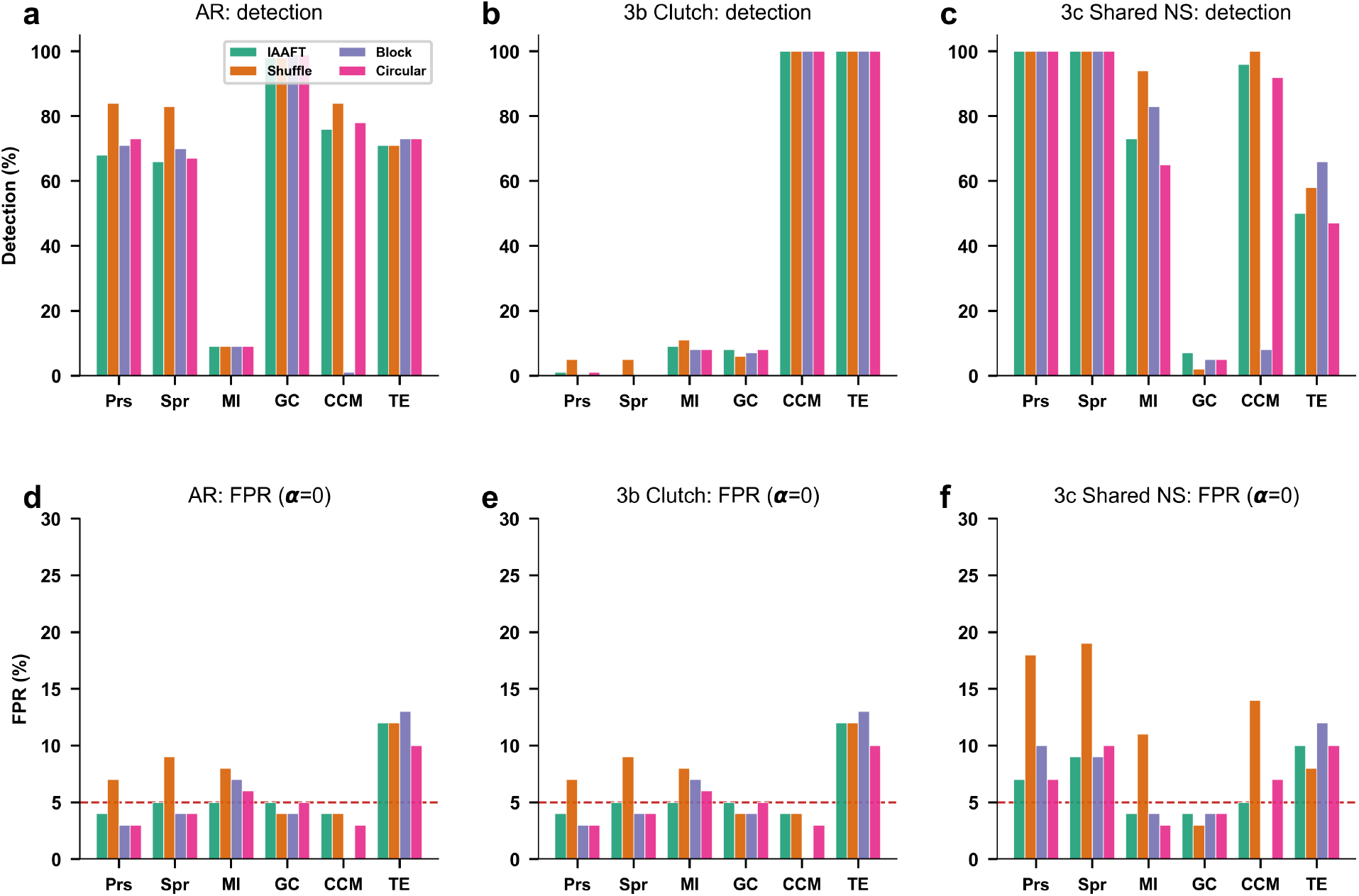
Surrogate choice across all six methods. Detection and false-positive rates for each method under four surrogate types (IAAFT, shuffle, block bootstrap, circular shift), extending the CCM-only comparison of Fig 3g,h. (**a–c**) Detection rate at each system’s swept coupling for AR (*α* = 0.30), biphasic clutch (*α* = 1.20), and shared non-stationarity (*α* = 0.10). Block bootstrap selectively collapses CCM (a, c) while other methods are unaffected. (**d– f** ) False-positive rate at *α* = 0 (no coupling); dashed line marks the 5% level. On the shared non-stationarity null (f), shuffle surrogates inflate false positives across the autocorrelation-sensitive methods, whereas IAAFT and circular surrogates remain near nominal. 1000 surrogates, 100 pairs per system.

